# Extracting physical characteristics of higher-order chromatin structures from 3D image data

**DOI:** 10.1101/2022.03.16.484676

**Authors:** William Franz Lamberti, Chongzhi Zang

## Abstract

Higher-order chromatin structures have functional impacts on gene regulation and cell identity determination. Using high-throughput sequencing (HTS)-based methods like Hi-C, active or inactive compartments and open or closed topologically associating domain (TAD) structures can be identified on a cell population level. Recently developed high-resolution three-dimensional (3D) molecular imaging techniques such as 3D electron microscopy with in situ hybridization (3D-EMSIH) and 3D structured illumination microscopy (3D-SIM) enable direct detection of physical representations of chromatin structures in a single cell. However, computational analysis of 3D image data with explainability and interpretability on functional characteristics of chromatin structures is still challenging. We developed Extracting Physical-Characteristics from Images of Chromatin Structures (EPICS), a machine-learning based computational method for processing high-resolution chromatin 3D image data. Using EPICS on images produced by 3D-EMISH or 3D-SIM techniques, we generated more direct 3D representations of higher-order chromatin structures, identified major chromatin domains, and determined the open or closed status of each domain. We identified several high-contributing features from the model as the major physical characteristics that define the open or closed chromatin domains, demonstrating the explainability and interpretability of EPICS. EPICS can be applied to the analysis of other high-resolution 3D molecular imaging data for spatial genomics studies. The R and Python codes of EPICS are available at https://github.com/zang-lab/epics.

## 1 Introduction

The genomic DNA is packaged into chromatin in a hierarchical structure in the eukaryotic cell nucleus. The higher-order chromatin structure has major impacts on gene regulation, cell identity, and human health [1, 2]. The fundamental unit of these structures are nucleosomes, DNA wrapped around a histone octamer core. The string of nucleosomes form various dynamic structures that can be characterized into domains such as topologically associating domains (TADs)[3]. Higher in the hierarchy, TADs can be categorized into compartment structures which can be defined as active (open) or inactive (closed). Open and closed chromatin have important biological roles. For instance, there is evidence that subtypes of cancers have different gene expression patterns associated with open and closed TADs [4, 5]. The structural properties of genome organization can be measured using high-throughput sequencing (HTS)-based technologies, like Hi-C [6, 7] and ChIA-PET [8], which primarily use DNA sequences as a positioning and quantification tool to determine the average proximity between two regions in the genome accumulated from a population of cells [6, 9]. Although the 3D configuration of chromatin can be inferred from Hi-C data using statistical or computational models [10, 11], direct measurement for physical characteristics of 3D chromatin structure in a single cell is still challenging [12].

Recent development of spatial omics technologies enables high-resolution detection of spatial distributions of molecular genomic information such as gene expression and genome organization by applying HTS techniques, such as 10X Visium [13] and Slide-seq [14], or using superresolution microscopy techniques with fluorescence in situ hybridization (FISH), such as seqFISH [15] and MERFISH [16]. Super-resolution microscopy has also been applied to measure chromatin structures directly, with emerged techniques such as 3D assay for transposase-accessible chromatin-photoactivated localization microscopy (ATAC-PALM)[17], 3D electron microscopy with in situ hybridization (3D-EMISH)[12], and 3D structured illumination microscopy (3D-SIM)[18]. ATAC-PALM is able to image the accessible genome at the nanometer scale and in conjunction with FISH-based techniques. 3D-EMISH is able to extract structures of probed genomic regions of interest and describe the domain structures at the nanometer scale. 3D-SIM is able to visualize chromatin throughout the cell with a 39.5 nm resolution. Furthermore, FISH-based chromatin imaging techniques such as ORCA [1] and MINA [19] have generated higher-order chromatin structures including TADs and A/B compartments consistent with what have been inferred from Hi-C, suggesting that chromatin domains (CDs) reconstructed from image data should reveal the same biological functions [18]. Compared with Hi-C, imaging-based methods provide more direct measurements of physical representations of higher-order chromatin structures directly in a single cell [12]. However, computational analysis of 3D chromatin image data remains a challenge. Specifically, computational models to connect active or inactive chromatin domains with the physical characteristics from 3D images are essentially nonexistent. Additionally, these computational approaches need to be developed while considering explainability and interpretability [20], so that biological insights can be generated from computational studies. Creating a computational model where scientists cannot ascertain the biological meaning is less useful than a computational model which can provide these insights. Thus, utilizing a machine learning (ML) approach which is easily interpretable and explainable is key for understanding the complex nature of 3D image data of chromatin.

In recent years, complex computational systems and advanced ML-based artificial intelligence (AI) have penetrated numerous fields of study [21]. Explainable AI (XAI) aims to provide explainable and interpretable insights to scientific inquiries [20, 22, 23]. Using XAI in biology means that the models and parameters can describe biological phenomena and characterize biological entities of interest in an explainable and interpretable fashion. Ensuring that the tenants of explainability and interpretability are met allows for scientists to evaluate if a computational model is adding to scientific knowledge. This work is built upon our previous model development work for extracting shape features from 2D image data such as blood cells[24] and satellite image classification [25], which adheres to the concepts of explainability and interpretability. Following the same principles, we extend our work for modeling 3D image data focusing on chromatin structure and to characterize 3D chromatin domains as open (active) and closed (inactive). This method allows for scientists to provide clear biological insights to various and dynamic structures of chromatin and their physical characteristics.

In this paper, we present Extracting Physical-Characteristics from Images of Chromatin Structures (EPICS), a method able to characterize different CDs from 3D image data generated from two techniques: 3D-EMISH and 3D-SIM. We apply EPICS to 3D-EMISH and 3D-SIM image data to characterize open or closed CDs from each data type. We then interpret the results of EPICS by identifying the most important physical characteristics as features that distinguish CDs for biological insights. Our work provides a spatial and physical perspective of 3D image data modeling for functional genomics.

## 2 Materials and Methods

EPICS is a computational method we developed to characterize CDs as open or closed from 3D image data. In this section, we first describe how EPICS reconstructs the chromatin from the raw image data. We then discuss our computational algorithm to determine if a CD is open or closed. This involves the selection of candidate metrics which are useful for determining if a CD is open or closed. It also involves identifying those variables which are the most important for classifying CDs from one another. EPICS is summarized in Figure 1a.

**Figure 1:**
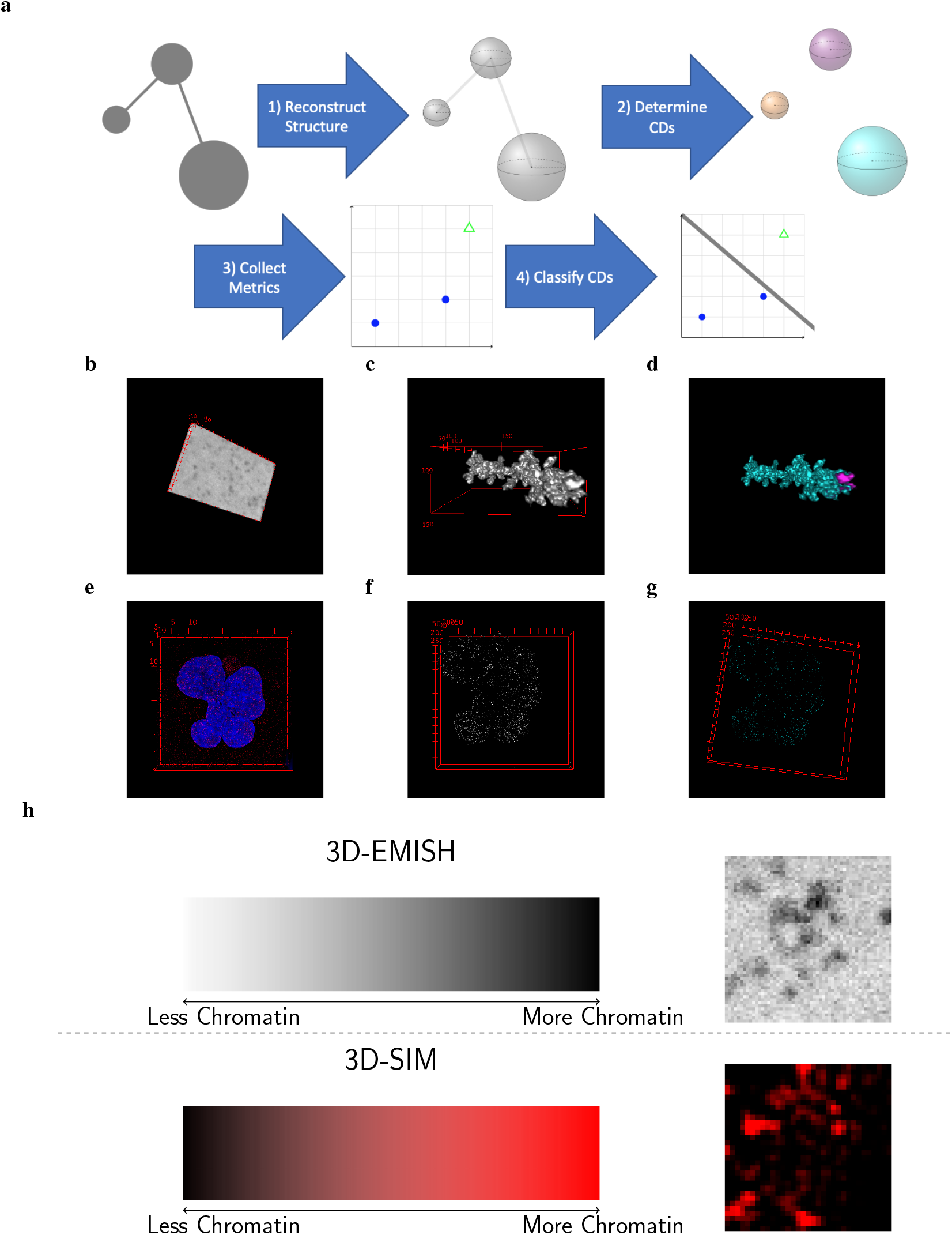
Schematic of EPICS with examples of images. **(a)** Schematic of EPICS. The gray circles represent the raw imaging data’s potential chromatin domains (CDs). The gray spheres represent reconstructed CDs. Each colorized sphere represents a uniquely identified CD (three in this case). The blue dots and green open triangle represent the closed and open CDs in the feature space. The line separating the points in the feature space represents the created model to classify the closed and open CDs from one another. The steps in **(a)** are exemplified with 3D-EMISH data in **b** - **d** and the 3D-SIM data in **e** - **g**. **(b)** Example of the raw input image from 3D-EMISH. **(c)** The reconstructed structure of the chromatin object of interest. **(d)** The resulting CDs. Each color represents a unique CD. In this case, there are two CDs with a larger cyan CD and a smaller pink CD. **(e)** The DAPI and H3K27me3 3D-SIM raw image data for the 30 hour treatment. **(f)** The reconstructed structure of the chromatin object of interest. **(g)** All identified CDs from the 30 hour treatment image. Each shade of cyan represents a unique CD. In this case, there are hundreds of different CDs present. The animated `.gif` files of **b** - **g** are provided at our GitHub link. **(h)** A color bar for each technologies raw pixel values. Examples of a cropped section of a slice from a 3D image is provided next to each graphical description.

### 2.1 Defining Chromatin Domain Assignment for Images

We describe our solution for reconstructing the chromatin domain structure from the raw image data while adhering to the concepts of explainability and interpretability herein. Using image operator notation to represent the image processing operators applied to the input data [26], we first smooth the image via

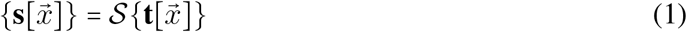

where 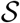 is the smoothing operator, 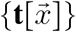 is the input image, and 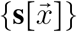 is the resulting smoothed image. We then isolate the relevant signals using

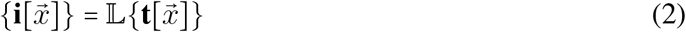

where 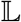 identifies the relevant signals of interest and 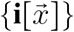 is the set of images containing the isolated signals of interest. We then apply

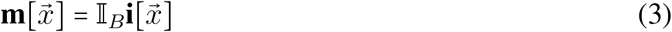

where 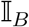 interpolates the object using the other slices to construct the missing slices and 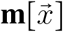 is the reconstructed object of interest from the given target signal image, 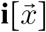 (Fig. 1a, Step 1). Interpolation is necessary to ensure that each voxel is approximately a cube in physical space. We then determine the CDs by

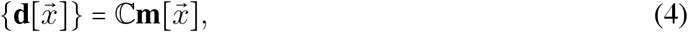

where 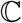 determines the CDs from the input image and 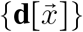 are the set of resulting CDs. The number of CDs is then determined (Fig. 1a, Step 2). The CDs are extracted from the 3D-EMISH and 3D-SIM data using Equations A32 - A35 and A39 - A40 respectively in the Supplementary Data.

We then collect a variety of explainable and interpretable metrics using the shorthand operator of 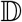 (Fig. 1a, Step 3):

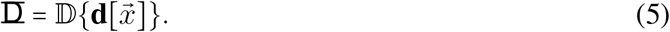

The equations for extract each metric are described in the Supplementary Data in Equations A13 - A27. From these extracted metrics in our resulting matrix, 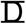, we build a model that creates rules for predicting whether a particular CD is open or closed. In other words,

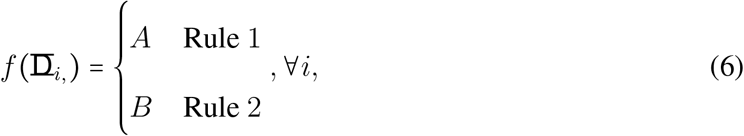

where *i* is the *i*^th^ CD such that *i* ∈ {1, …, *N*}, *N* is the total number of CDs, and *A* and *B* are the two possible chromatin states, consistent with A/B compartments inferred from Hi-C data. A is considered active or open, while B is inactive or closed (Fig. 1a, Step 4).

This computational approach is similar in spirit to the computation required for A/B compartment assignment using data from Hi-C. EPICS and the computation using Hi-C data both use a set of rules to determine if a target is open (active) or closed (inactive). While Hi-C is based on correlations, EPICS uses a logistic regression model based on the physical characteristics of the objects. Supplementary Data A.1 provides an overview of the computation for A/B compartment assignment using Hi-C data.

### 2.2 Defining Open and Closed Domains for Images

We need to identify potential candidate metrics that would be useful for clustering the CDs and classifying the identified CDs. To that end, we need to explicitly state what characterizes open and closed domains. Open domains have lots of space in between points and will be sparser. Closed domains are compact and close to one another. Natural choices for capturing these sparse and dense domains would be shape-based and intensity-based metrics. Shape metrics describe how dense the domain is spatially as indicated by Figure B1. The intensity metrics describe the amount of the chromatin present in the sample. For 3D-EMISH data, low values indicate more object of interest, while larger values indicate that less object of interest is present (Fig. 1h). To this end, we select 8 shape and 11 intensity metrics to describe our domains of interest, as summarized in Table 1.

**Table 1:**
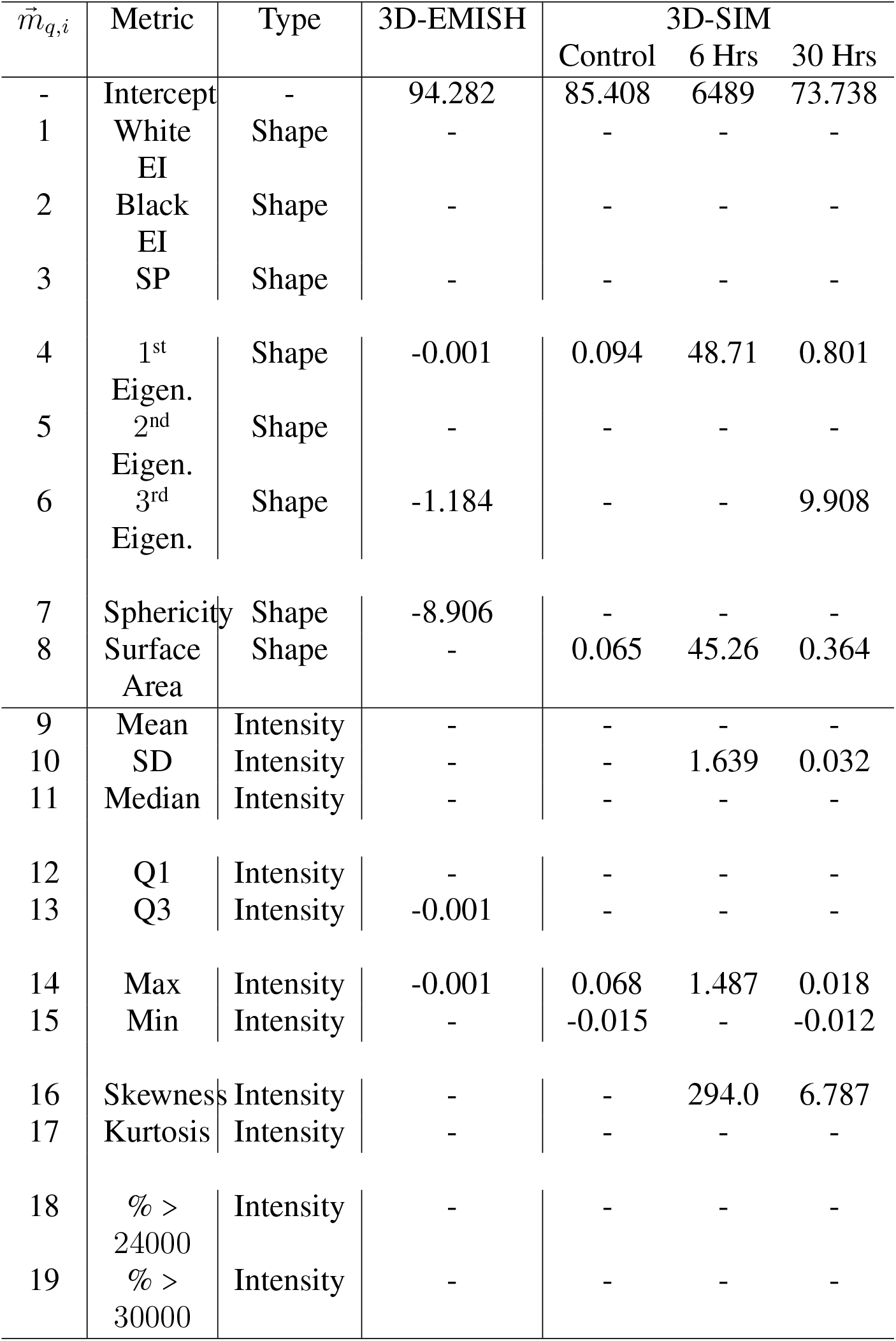
Metrics used in EPICS on a given CD, *i*. The first column is the *q*^th^ metric, where *q* ∈ {1, 2, …, 19}. The last columns correspond to the final estimated parameter for the logistic regression (LR) models for each data sets. refers to features not included in the model. 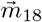 and 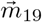 are not used for 3D-SIM data.

The first shape metrics are the SPs and EIs, which are collected by extending shape proportion and encircled image-histogram (SPEI) algorithm [27] to be applicable to 3D shapes. The EI is the black and white pixel counts of the shape after the shape is placed in the minimum encompassing sphere and then the minimum encompassing cube. In other words, this is the volume and the surrounding volume of the object of interest. The SP value is the proportion of the volume of the shape relative to the sum of the EI. The other shape metrics collected that are used in the model are the eigenvalues of the shapes [26], sphericity [28], and surface area. The eigenvalues measure the major and minor axes of the shape. Sphericity measures how spherical a given shape is. Surface area is the 3D perimeter of the object of interest. This results in a total of 8 total shape metrics.

The intensity-based metrics are merely their respective statistic for the object’s intensity based values. For example, the mean of the intensity values measures the arithmetic mean of only the object’s voxel values. This does not include the background of the object. For the Mean, Median, Q1, Q3, Max, and Min Intensity metrics, low values indicate more or less chromatin for 3D-EMISH or 3D-SIM, respectively. High values indicate less or more chromatin for 3D-EMISH or 3D-SIM, respectively (Fig. 1h). The remaining statistics are interpreted and explained in the typical manner. There are a total of 11 intensity-based metrics.

Extended explanations, interpretations, and image operators for each shape and intensity metric are provided in the Supplementary Data. In short, each of the metrics provided is explainable and interpretable. This helps to make the results of describing the open and closed chromatin domains explicitly understood and aid in understanding their biological underpinnings.

### 2.3 Determining Rules for Open-Closed Chromatin Domains

Here we expand the notation from Equation 6 to provide an explicit description of the analysis done in EPICS. For notation purposes, we have *J* batches such that *j* ∈ {1, 2, …, *J*}. For the 3D-EMISH data, *J* = 2. For the 3D-SIM experiments, *J* = 1 since each experiment has a different treatment. Further, we have 19 features such that *q* ∈ {1, …, 19}. Thus, ∀*q*, *j*, we perform

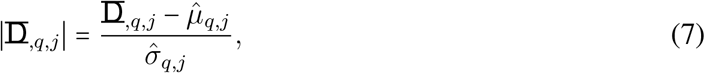

where 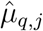 and 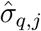 are the sample mean and standard deviation of the *q*^th^ feature for the *j*^th^ batch. We do this to normalize the data and remove batch effects. We then perform the following to obtain the estimated A/B chromatin states:

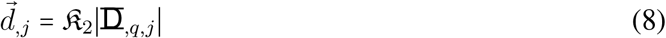

where 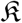 is the *k*–means clustering operator. In this case, we perform *k*-means clustering with a known number of clusters of 2. The output is the estimated chromatin states saved in an associated vector, 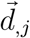. Here, we inspect each cluster and determine which cluster is associated with open and closed for each batch. These are the open and closed class labels to be further modeled in future steps of EPICS.

However, this is insufficient for determining clusters as we need to potentially interpret which cluster is open and which is closed. This introduces human subjectivity and errors into the analysis. Further, the *k*-means clustering model does not provide any meaningful insight to which are the most important variables for discriminating the open and closed CDs from one another. Thus, we select a logistic regression (LR) model to describe open and closed CDs [29]. However, there are too many variables for this to be a truly interpretable and explainable model [20]. Thus, we use the least absolute shrinkage and selection operator (LASSO) algorithm to select the most important variables for our model[30, 31, 32]. Thus, we first model:

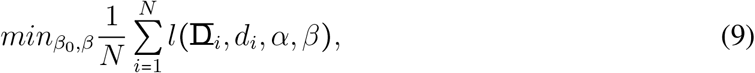

subject to ||*β*||_1_ ≤ λ where ||·||_1_ is the *L*_1_-norm and λ is a tuning parameter [32]. The data was split into the training and validation data using a 70-30 split. We ensured that the proportions of the open and closed CDs were preserved in the training and validation data using stratification[33]. We select the tuning parameter, λ, by using 10-fold cross validation on the training data. While the optimal λ is able to obtain a very high classification rate, it retains a larger number of variables. Thus, we select the λ value within 1 standard error for the 3D-EMISH data since it also has a very high classification rate, has less variables retained in the models, and is the more prudent choice (Supplementary Fig. B2). After repeating this process for the 3D-SIM, we chose the same value for the 3D-SIM’s choice of λ as the 3D-EMISH’s λ value to be more conservative [30, 31] (Supplementary Figs B3–B5). Extended discussions on the choice of λ for the 3D-SIM data are provided in the Supplementary Data.

After we obtain the non-zero coefficients, we use those variables to model the following:

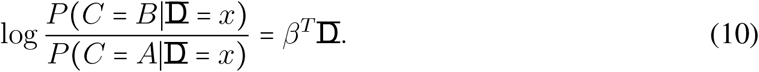

The final estimates of the variables’ coefficients are found in the third through sixth columns of Table 1.

The parameters of the LR model are typically described using the log-odds ratio or the odds ratio [34, 29]. Thus we are able to interpret the log-odds ratio in the following manner for the *q*^th^ variable: assuming that all of the other variables are held constant, for every one unit increase for the given variable, we expect the log-odds of a being a closed CD to increase by 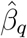 [34]. Positive values for estimated coefficients, 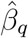, and an increase in the associated variable corresponds to increasing the probability of being closed. Conversely, negative values for the estimated coefficient and an increase in the associated variable would indicate a decrease in the probability of being closed. Further, we are able to convert these to odds by taking the exponential of the estimated coefficient value.

For example, if we assume that all other variables do not change for the 3D-EMISH model, for every unit increase in the maximum of the intensity value of the CD, we expect the log-odds of being a closed CD to decrease by 0.001. The odds would be *e*^−0.001^ = 0.999. Conversely, for the 3D-SIM image with the 6 hour treatment, we would expect the log-odds of a CD being closed to increase by 0.068. The odds would be *e*^0.068^ = 1.070. This can be repeated for each variable of interest for each model.

This allows analysts to compare and contrast different models. Further, we are able to identify the similarities and differences between different data and treatments. For example, the 3D-SIM Control and 6 hours identify different parameters as being the most important. This suggests that there are tangible differences between the CDs. In other words, we expect that the 6 hour treatment is fundamentally changing the CDs. To quantify these difference, we can observe the parameter differences between the two models. For example, the max intensity parameter value differs by a factor of 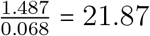. Thus, the 6 hour treatment impacts the max parameter value by about a factor of 21. Thus, this provides additional evidence that the treatment changes how CDs exist in physical space.

Our EPICS method is considered XAI since they use features that are both interpretable and explainable: the LR model is a highly interpretable and explainable model to discriminate open and closed CDs[20], and each operation performed during the entire process is clearly described while also being interpretable and explainable. The reduction in complexity is exemplified in Figure B6.

### 2.4 Materials

Raw 3D-EMISH data was obtained from Trzaskoma *et. al*.’s GitHub (https://github.com/3DEMISH/3D-EMISH)[12]. Raw 3D-SIM images were obtained from Cremer et. al.’s paper[18]. The computation was performed on a Ubuntu 18.04.5 system with 64 GB of RAM and an Intel® Xeon(R) W-2245 CPU @ 3.90GHz with 8 cores and 16 threads. For the image processing and metric collection, we used Python 3.6.9[35] alongside numpy[36], scipy[37], skimage[38], sklearn[39], and kneed[40]. Determining the CD assignment was performed in R[41] using the xtable[42], caret[43], glmnet[44], and clue[45] packages.

## 3 Results

We applied EPICS on 3D chromatin image data from two techniques, 3D-EMISH (Fig. 1b - 1d, Supplementary Figs. B7–B12) and 3D-SIM (Fig. 1e-1g, Supplementary Figs. B13–B16). We identified open and closed chromatin domains (CDs) from each data type, and explained and interpreted the model and results by extracting important physical characteristics from the image data. In addition, we identified batch effects from 3D-EMISH data and demonstrated that EPICS is able to characterize open and closed CDs despite the batch effects.

### 3.1 Determination of Open or Closed Chromatin Domains from 3D-EMISH data

The 3D-EMISH data for our analysis is from Trazaskoma et. αl.[12]. They analyzed human lymphoblastoid cells by probing a 1.7 mega-base (Mb) segment of the genome and extracted 229 image stacks or z-stacks of potential targets. Each voxel intensity value corresponds to a measured object as summarized in Figure 1h. White intensities indicate less material, while darker colors correspond to more. Our computational analysis using EPICS determined out of 451 CDs extracted from the 229 images, there exist 163 and 288 closed and open CDs, respectively, across the two experiments. 19 shape and intensity-based measures of CDs were used to determine these classes using *k*-means clustering. We then used LASSO to select the appropriate features that are able to discriminate between the open and closed CDs across the different batches (Figure 2).

**Figure 2:**
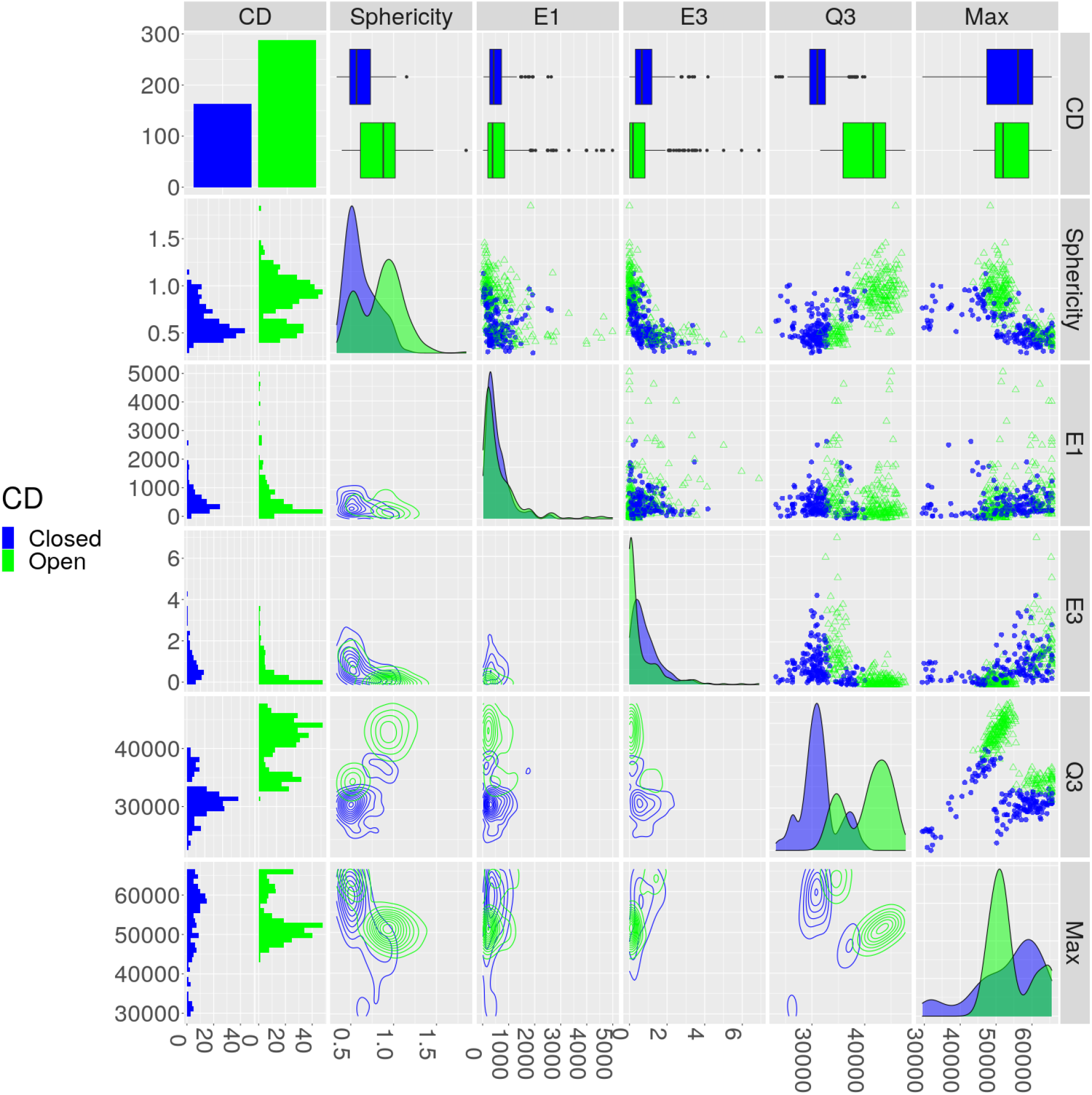
The important features for discriminating between open and closed CDs from the 3D-EMISH data. Open and closed CDs are green and blue dots, respectively. The top left cell provides the counts of closed and open CDs. The remaining cells in the top row provides a boxplot of each feature. The remaining first column provides a histogram of the closed and open CDs for each feature. The remaining diagonal cells provide the density plots of each feature by each class. The remaining upper triangular portion and bottom triangular portion provide the 2D scatterplots and the 2D contour plots, with corresponding axes labeled for each column and each row.

We then sorted these five variable from the most to least important using Variable Importance (VI). VI is the absolute value of each variables associated z-value from the LR model. Relative VI is VI divided by the largest z-value. The 3^rd^ quantile (Q3) is the most important variable. The intensity-based metrics are the two most important variables, while the shape-based metrics are the three least important (Fig. 3a, Supplementary Table C1). Using the top three variables provides a clear separation in 3D space (Fig. 3b).

**Figure 3:**
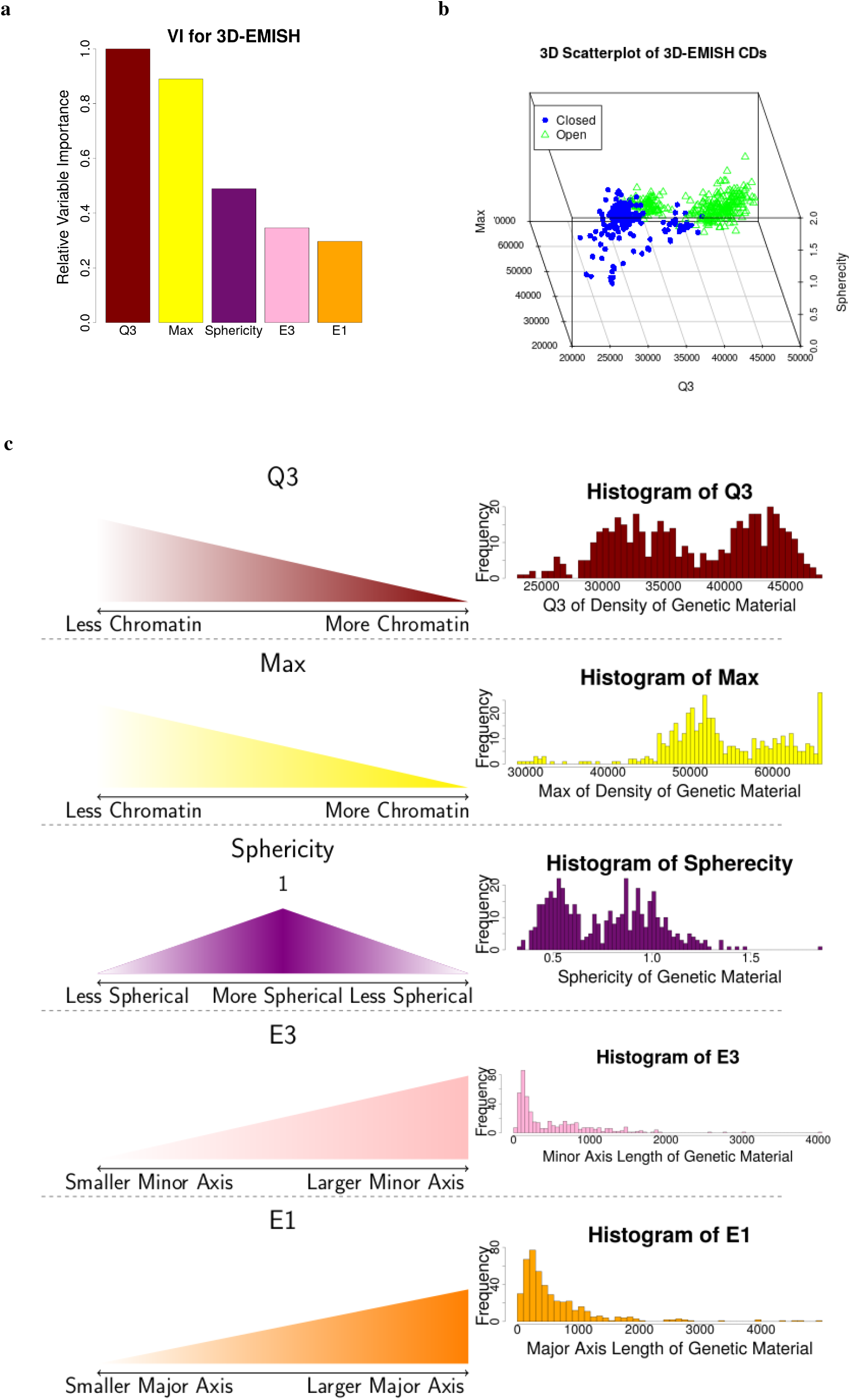
EPICS identified important biological features of CDs from 3D-EMISH images. **(a)** Relative variable importance (VI) of the 5 most important features from the 19 candidate features. **(b)** The 3 most important variables in a 3D scatterplot. **(c)** Value interpretation and distribution of each identified important features for the 3D-EMISH data.

We identified five most important features to discriminate open and closed CDs. Closed CDs tend to have lower intensity values, as showcased by Q3. These lower intensity values indicate that more chromatin is present. This means that lower intensity values correspond to a denser object. Conversely, higher intensity values indicate that less chromatin is present. Thus, higher intensity values indicate that chromatin is more sparse or spread out. The maximum of the intensity of closed CDs are smaller than those that are open. Thus, closed CDs have more chromatin and are more densely packed than open CDs. The sphericity of open CDs tend to be closer to 1 than closed CDs. Thus, open CDs tend to be closer to a sphere in shape compared to closed CDs. There are no obvious patterns for differentiating between open and closed CDs when using the relative major and minor axis lengths (E1 and E3, respectively). However, alongside other metrics, their impact becomes more apparent (Fig. 2). All of these features have graphical representations provided in Figure 3c. Thus, open CDs tend to have less chromatin material and tend to be more spherical in shape, while closed CDs tend to have more chromatin material and are less spherical in shape.

The logistic regression (LR) model created to discriminate open and closed CDs using the 3D-EMISH data across the experiments was able to achieve an overall classification rate of about 95% (Table C2) and 96% (Table C3) on the training and validation data, respectively. Their associated 95% confidence intervals (CIs) for the overall classification rate were (92%, 97%) and (91%, 98%), respectively. Thus, our classification model performs well on data not used to build the model [20]. Further, since we used a LR model, our solution is highly interpretable and explainable [20].

### 3.2 Batch Effects exist in 3D-EMISH data

Our analysis shows that strong batch effects exist for 3D-EMISH data. The first 3D-EMISH experiment published in Trazaskoma *et. al*. [12] was 7 × 7 × 30 nm for each voxel, while the second experiment was 5 × 5 × 30 nm. After correcting for the differences in voxel resolutions during the reconstruction step, we observed that CDs identified from the two batches can be clearly separated in the feature scatterplot matrix (Fig 4a), indicating strong evidence of a batch effect. We confirmed this by building a LR model to predict the batch using the six variables identified by LASSO. This model was able to achieve about 98% and 99% overall accuracy on the training and validation data, respectively. Thus, future analyses should account for different voxel resolutions.

**Figure 4:**
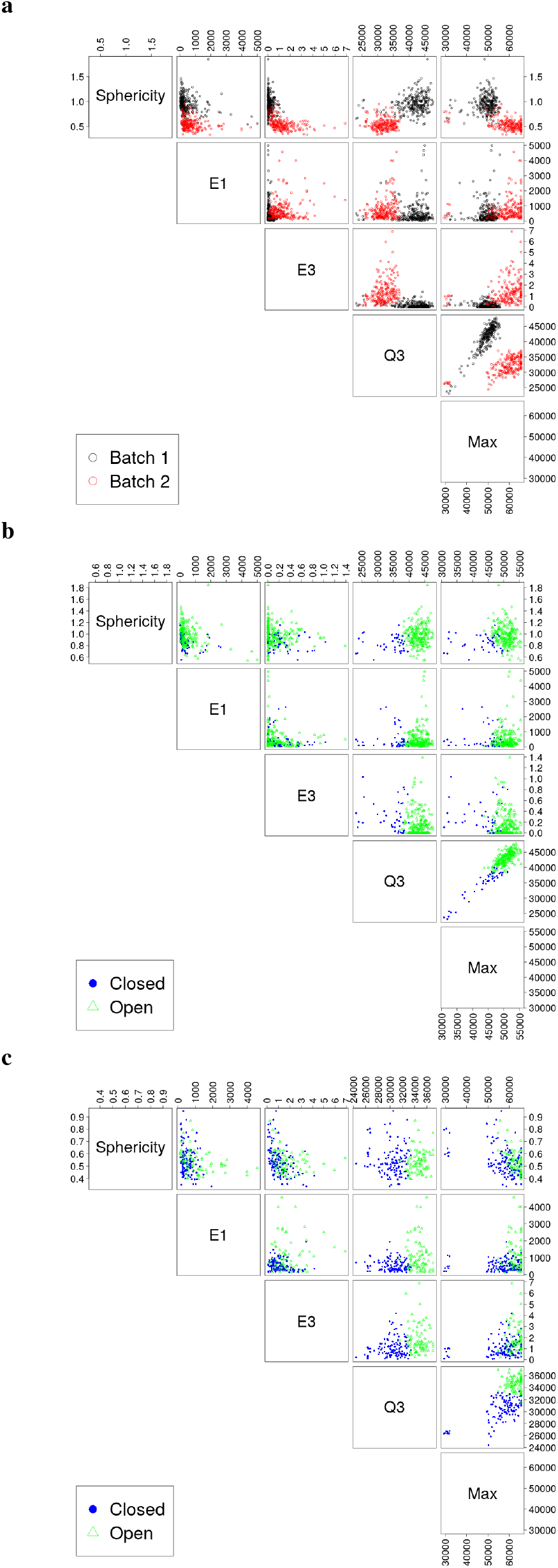
EPICS is able to account for batch effects in their original space. **(a)** Scatterplot matrix showing the features from Figure 2, but with colorizations corresponding to the first and second experiments. The black and red open circles correspond to the first and second experiment, respectively. **(b, c)** Scatterplot matrices showing the CDs from the first experiment **(b)** and from the second experiment **(c)**.

To account for the batch effect, we used EPICS to first cluster per each batch using batch removal and *k*-means independently before combining the data for the LASSO. Batch removal was performed by subtracting the sample mean and standard deviation of for each respective feature by batch (Eq. 7). Typical batch removal requires the analysis to remain in an abstract space. However, we found that EPICS is still able to classify the open and closed CDs in the original feature space (Figs 4b & 4c). For instance, the blue solid dots are left of the green open triangles across batches for Q3 against Sphericity, E1, or E3. While the two experiments have different ranges for the features selected by LASSO (Figs 4b & 4c), EPICS is able to use the clusters found from the independent batch removal step in the LR model using the original physical space for the variables (Figs. 2 and 3b). Thus, EPICS is still able to capture the important biological features across batches in their original physical space and identify open or closed CDs.

### 3.3 EPICS identifies chromatin domains from 3D-SIM data

To test the general usability of EPICS, we applied EPICS to 3D image data generated from other techniques. Specifically, we applied EPICS to 3D-SIM generated immunostained images for repressive histone modification H3K27me3 in human colon cancer cell lines [18]. We identified open and closed CDs of the cell under three different conditions: control, treated for 6 hours in auxin, and treated for 30 hours in auxin (Figure 5, Supplementary Tables C4–C6). Each dataset was an image stack. Each voxel intensity value corresponds to a biological object as summarized in Figure 1h. Large intensity values indicate more chromatin, while smaller values correspond to less. We processed the 3D-SIM data with the similar procedure for 3D-EMISH, as shown in parallel in Figure 1. For example, we identified the important variables and characterized the physical properties of the H3K27me3-marked chromatin in these cancer cells under different treatments (Supplementary Tables C4–C6), and provided overall accuracy measures and confidence intervals (CIs) for each model’s performance (Supplementary Tabels C7 - C12). The smallest accuracy on the validation data was about 0.990 with an associated 95% CI of (0.980, 0.995).

**Figure 5:**
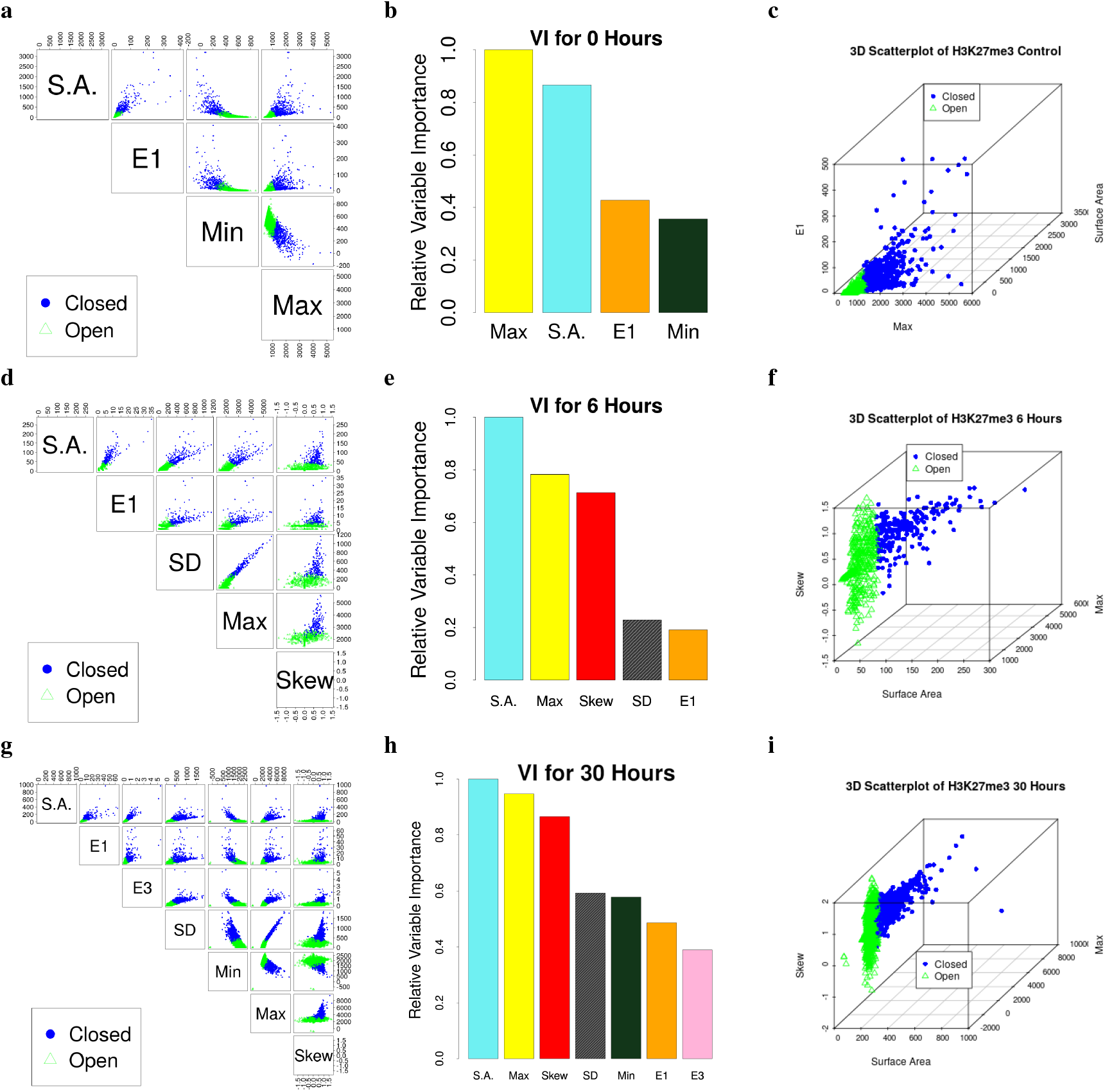
EPICS identified open and closed CDs from 3D-SIM data for H3K27me3 and a human colon cancer line across different treatments. Open and closed CDs are presented using green open triangles and blue closed circles, respectively. S.A., Surface Area. **(a, d, g)** Scatterplot showing the EPICS-identified important variables. **(b, e, h)** The relative variable importance (VI). **(c, f, i)** A 3D scatterplot of the top 3 most important variables clearly separating open and closed CDs for a given sample.

There are two primary biological insights obtained from these results. The first is the nature of open and closed CDs at a coarser resolution using 3D-SIM relative to the finer resolution of 3D-EMISH. When compared to open CDs, closed CDs tend to have more chromatin material, a larger surface area, and have very dense sections within the CD. Thus, closed domains appear as large, dense, asymmetric chromatin. When compared to closed CDs, open CDs tend to have less chromatin material, less surface area, and a smaller major axis. Thus, open domains tend to be small, sparse, symmetric chromatin. The second biological insight is the increase in complexity required to classify open and closed CDs across treatment types. There is evidence that CDs exist across different treatments [18]. However, our models indicate that the CDs are not static and remain unchanged across treatments. In fact, our model shows that the treatments might change the physical characteristics of the chromatin. In particular, since the 3D-SIM data has three treatments, we can quantify the differences between models for classifying the open and closed CDs. For example, each of the 3D-SIM models retained surface area and the max intensity. However, the parameter values in our models are drastically different from one another. Such differences might be due to technical variances on the data or the model, or could be real biological results. Further investigation is needed on the EPICS results from 3D-SIM to provide insights to the biological nature of chromatin.

## 4 Discussion

In this work, we present a computational method, EPICS, which we developed to identify chromatin domains from 3D image data and to determine if identified chromatin domains are open or closed based on its physical representation from the images. We used data generated from 3D-EMISH and 3D-SIM techniques as two case studies to exemplify the workflow, functions, and results of EPICS, as well as the ability to correct for batch effects for 3D-EMISH data. More importantly, we demonstrated the ability of EPICS to explain and interpret the model and results. This allows researchers to learn the physical characteristics from super-resolution imaging data to understand the morphological properties of higher-order chromatin structures such as open and closed chromatin domains or active and inactive compartment structures. From the 3D-SIM data in cells under different treatments, we provided evidence that open and closed chromatin domains’ physical characteristics might dynamically change in time under treatments. Furthermore, we incorporated the tenants of explainability and interpretability into the development of EPICS to ensure the ML methods used are within the confines of XAI. To our knowledge, EPICS is the first and so far the only XAI method for analyzing 3D chromatin image data.

Specifically, we showed that EPICS is able to characterize open and closed CDs from 3D-EMISH images at resolutions as small as 5 × 5 × 30 nm and 3D-SIM images at 39.5 × 39.5 × 125 nm. We identified the variables that were the most important to discriminate open and closed CDs for each of our models and data sources. The LR model was able to achieve an overall classification rate of about 94% on the 3D-EMISH validation data, which were not used to build the model. The 3D-SIM model was able to achieve accuracies of at least 98% on the validation data. Thus, the models should generalize to outside data well. Therefore, we provide a computational framework that is able to describe and classify open and closed CDs from these two different technologies.

Logistic regression (LR) is highly explainable since we can describe the exact relationship modeled and how the parameters of the model work. LR is interpretable since each parameter corresponds to a physical change in the probability of belonging to a particular class, and the final LR parameter values are shown in Table 1. Thus, LR is a prime example of a model which is able to satisfy the conditions of an XAI model.

EPICS handles the batch correction by clustering the data by batch before the model is built. This is vital as these CDs were collected over two different datasets, but EPICS is able to classify the clusters using the LASSO and LR in their original unchanged space. The final LR model enables future analysts the ability to predict new observations without needing to do any batch correction for 3D-EMISH data.

Lastly, EPICS reduces model complexity by identifying the important features from the image data to consider as exemplified in Figure B6. By removing less important variables, biologists are able to make more insightful and meaningful inferences. By using the LASSO, we are able to identify the relevant features for understanding the underlying biological properties of the CDs. Thus, EPICS allows analysts to interpret the results in the original and physical units of the collected features.

This is the first work to characterize higher-order chromatin structures from image data in this manner. Chromatin structures from 3D-SIM data across different treatment for cells using volume were found to have statistically significant differences between the treatments [18]. Others used sphericity, surface area, and volume to find statistically significant differences between chromatin structures that have different number of domains [12]. However, none of these approaches use a large number of features to describe the physical characteristics of chromatin structures, nor do they use these features to identify clusters of open and closed CDs. Thus, EPICS is the first approach to identify open and closed CDs using the physical characteristics of chromatin.

The identification of open and closed CDs from 3D chromatin image data using EPICS is analogous to the identification of A/B compartments from Hi-C data. We speculate that they refer to comparable structural information of chromatin states. However, due to lack of orthogonal information such as genomic coordinates for validation, the ground truth of what open/closed CDs actually mean remains unclear. Further studies are needed for finding more biologically meaningful interpretation of these computationally-determined chromatin states.

There are three primary pitfalls of EPICS. The first is that some steps in the image pre-processing differ between the two technologies. This is primarily due to the nature of the different types of raw data. However, further improving the similarities in the algorithm would help ensure that EPICS treats the different raw data from different technologies as equitably as possible. The second is the human task of interpreting the results from *k*-means clustering. To that end, some of this could be alleviated by automating some of these cluster assignments in future versions of EPICS. The third is the lack of genomic coordinate information in the two types of image data presented in this work. This limits the possibility to further interpret the CD classification result of EPICS for functional association with the genome, and presents us from using orthogonal information such as Hi-C data to validate our image-based domain inference. Potential application of EPICS to other FISH-based spatial genomics data that barcode genomic information, such as seqFISH, ORCA, or MINA, can further improve the interpretability of EPICS for functional genomics studies. Nevertheless, considering that these pitfalls can be managed and improved in future work, EPICS potentially has a broad application in image-based spatial genomics data analysis.

## 5 Conclusion

EPICS allows us to understand the biological underpinnings of cells at a new level for investigation. Using our model for classifying CDs from 3D-EMISH data or 3D-SIM would allow for deep understandings of a variety of chromatin structures. Unlike Hi-C, 3D-EMISH and 3D-SIM are able to provide physical representations of the chromatins. Further, 3D-EMISH and 3D-SIM provides a more direct measurement of chromatins than Hi-C. Thus, EPICS has the potential to yield new and meaningful biological insights for chromatin structures captured using 3D image data.

## Acknowledgements

This work was supported by the US National Institutes of Health [R35GM133712] to C.Z.

## Supplementary Data

### A Supplementary Text

#### A.1 Hi-C Compartment Assignment

We include a summary of the computation involved for Hi-C compartment assignment from Lieberman-Aiden *et. al*. [6]. We provided their framework here explicitly to precisely compare and contrast our approach to theirs. Their first step is to

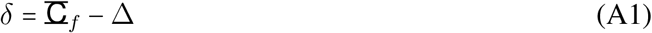

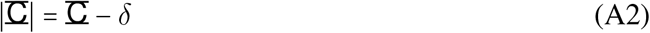

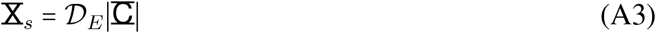

where Δ a 3 × *i* is the location of each Fiducial bead in each tensor, *δ* is the correction factor to be used for each image after MERFISH is applied, 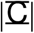 is the corrected distances, 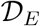 calculates the Euclidean distance between the compartments, and 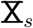 is the resulting *i* × *i* spatial distance matrix. They then estimate coefficients, 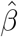, to obtain a normalized genetic distance matrix, 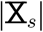, by performing:

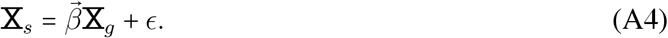

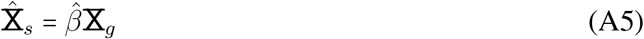

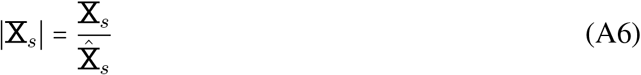

where 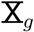 be the genetic distance *i* × *i* matrix. A genetic distance between two genes measure how similar a pair of genes and 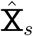 is the predicted *i* × *i* matrix. Since the Normalized matrix, 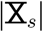, is not guaranteed to be 1, we need to perform these operations to guarantee that this occurs. Thus, they perform

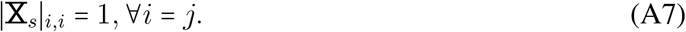

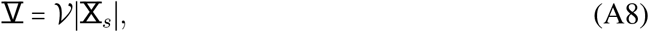

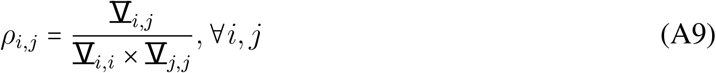

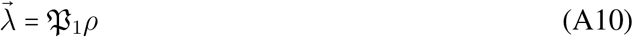

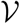 extracts the covariance matrix and 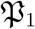 performs principal component analysis and extracts only the first dimension’s *i* coefficient values and outputs them to the vector 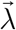. Lieberman-Aiden *et. al*.observed that the first eigenvector corresponds to compartment assignment[6]. Thus, they suggest that:

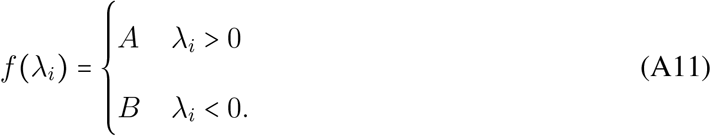

However, they make their most important inferences using a correlation matrix. Thus, they weaken their inferential capabilities by relying upon correlation[46, 47].

#### A.2 Simulating Chromatins Data Exploration

This paper describe the first analysis to use shape metrics to classify CDs as open and closed. To this end, we simulated genomic structures using multivariate normal or Gaussian distributions to ensure that using shape metrics would be reasonable. The two multivariate normal distributions that were selected that shared the same mean, *μ*. The covariance matrices of the first and second distribution were, respectively, 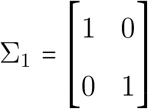, and 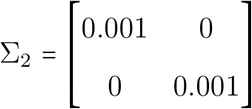. The first covariance matrix corresponds to open CDs, while the second corresponds to closed CDs. For each simulated structure, we randomly selected 1000 observations. Using image operator notation to represent an image processing algorithm[26], these two distributions are then converted into 2D images using

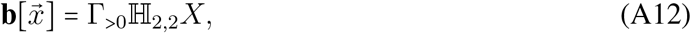

where *X* is the input data, 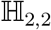 converts the data into a 2D histogram and Γ_>0_ is the threshold image operator. This equation converts numeric data of the sparse and dense simulated structures into image like those provided in Figures B1a and B1b, respectively. This is then repeated 100 times to collect 100 images of these simulated structures for a total of 200 images. From each of these images, a variety of shape metrics are collected such as area. A sample of these shape metrics are provided in Figure B1c with the red dots representing the sparse structure and the black dots representing the dense structure. Since the two classes of structures are separable, we could easily create a model to classify these simulated genomic structures.

#### A.3 Metric Collection

##### A.3.1 Shape Metrics

We collected the encircled image-histograms (EIs) using the algorithm described by Lamberti [27]. This algorithm results in a vector 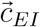 which contains the white and black pixel counts from image 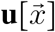. These counts are the first two metrics, 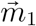 and 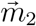, respectively. The shape proportion (SP) value for a given image, *i*, is merely

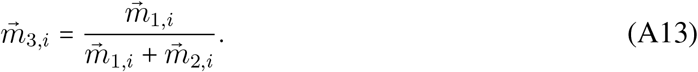

The SP value is essentially the proportion of white pixels after applying the SPEI algorithm [27]. This SPEI algorithm puts a shape in its minimum encompassing circle. Then the circle is placed in its minimum encompassing square. However, since our images are in 3D, we extended the SPEI algorithm in the manner suggested by Lamberti. He suggests to use a sphere in place of the circle and a cube in place of the square. The EIs are the white and black pixel counts after applying SPEI [27]. Lamberti states that the metrics from SPEI are highly explainable and interpretable[20].

To obtain the eigenvalues, we performed

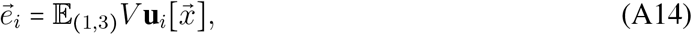

where *V* collects the covariance matrix of the shape matrix and 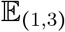 calculates the first, second, and third eigenvalues of the resulting covariance matrix. The *j*^th^ eigenvalue on image *i* is 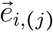.

The first, second, and third eigenvalues are metrics 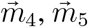, and 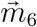, respectively. It is well-known that the eigenvalues of a covariance matrix correspond to the linear combination in the data which maximizes the variance for their respective dimension [48]. For instance, the first eigenvalue is the linear combination of the data which maximizes the first eigenvalue [48]. We also know that the linear projections, or eigenvalues, are orthogonal to one another [48]. Thus, the eigenvalues are measures of each of the three axes of our given shape. Lamberti states that the eigenvalues have a medium level of explainability and interpretability[20].

Sphericity, 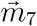, was collected on image *i* by

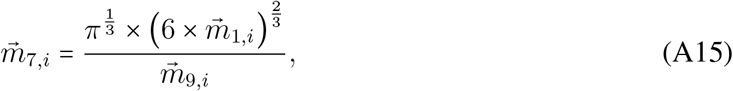

where 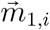 is the White EI or more commonly known as volume for 3D images and 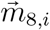 is surface area. Values close to 1 are more spherical. Values farther away from 1 are less like spheres in shape. Sphericity is very similar in spirit to circularity. The clear definition make sphericity highly explainable. However, sphericity is only somewhat interpretable. For example, sphericity does not provide guidance for what to classify the shape of a CD with a value of 2. The CD is certainly less spherical than a CD with a value of 1. However, sphericity is not equipped to answer interpret the metric with any physical meaning beyond being more or less like a sphere.

Surface area, 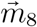, was collected on image *i* by

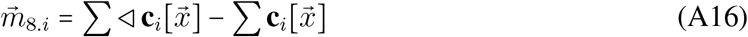

Surface are is both highly explainable and interpretable. It is explainable since we can accurately describe how to calculate it. It is interpretable since surface area describes the amount of space the CD’s outermost layer occupies.

##### A.3.2 Intensity Metrics

The features for intensity were the mean, standard deviation (SD), median, first quantile (Q1), third quantile (Q3), maximum, minimum, skewness, kurtosis, percentage greater than 24,000, and percentage greater than 30,0000 of the object’s pixel intensity values. The object’s pixels were extracted by using the segmented shape. Explicitly, for a given image and mask, metrics 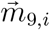 to 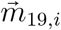

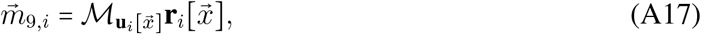

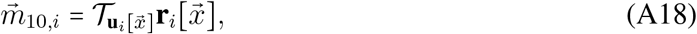

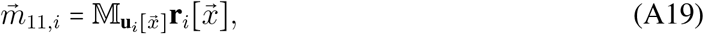

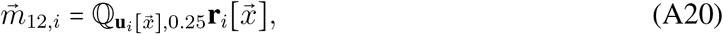

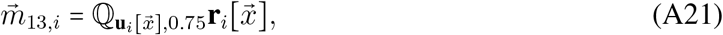

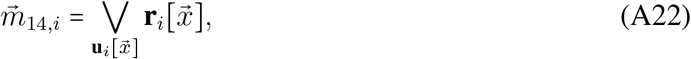

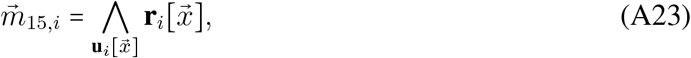

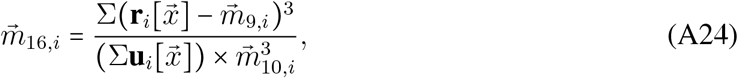

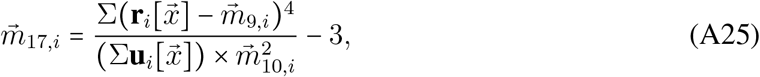

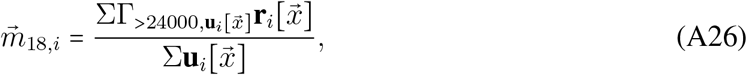

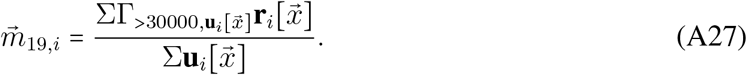

where 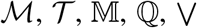, and 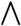 are the mean, standard deviation, median, quantile, max, and min operators, respectively. All of these operators are only extracted on the 1’s of image 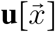, which ensures that only the colors of object are considered for a given input image. Note that a intensity value has different meanings across technologies. For example, small values for 3D-EMISH mean that more of the chromatin is present and larger values mean that there is more. Conversely, small values for 3D-SIM mean that there is less genetic material, while large values mean that there is more. This is summarized in Figure 1h The following paragraphs describe each metric in details. This will show how our intensity features have an interpretable and explainable meanings.

The mean value for a given channel indicates how much of that color is in the respective channel. For 3D-EMISH, large values indicate that target chromatin is not present in the image, while small correspond to the image having larger amounts of a given genetic target. The opposite is true for 3D-SIM. Larger values mean that there is more genetic material, and smaller values mean that there is less genetic material. Thus, the mean intensity value is explainable since it describes how it captures the typical amount of the genetic target. It is interpretable since the mean intensity value corresponds to physical definitions about the genetic target of interest. The median intensity value follows a similar logic to the mean.

The SD indicates how much the presence of a color changes throughout an image. A large SD indicates that the presence of the intensity of a color changes dramatically, while small a SD corresponds to a fairly uniform representation of the intensity of a color. Thus, the SD intensity value is explainable since it describes how it the amount of genetic target varies. It is interpretable since the SD intensity value corresponds to physical definitions regarding the variation of the genetic target of interest in a given object.

The Q1 and Q3 values indicate the amount of genetic material that separates the sample observation’s intensity distribution into the 25^th^ and 75^th^ quantiles, respectively. For 3D-EMISH, large intensity values indicate that target chromatin is not present in the image, while small correspond to the image having larger amounts of a given genetic target. For 3D-SIM, larger intensity values mean that there is more genetic material. Smaller values mean that there is less genetic material. Thus, the quantile values are explainable since it describes the dividing point of the sample distribution of an observation of the genetic target. It is interpretable since quantile values corresponds to physical definitions about the genetic target of interest.

The max and min values indicate the maximum and minimum intensity values observed in a given observation, respectively. For 3D-EMISH, large values indicate that target chromatin is not present in the image, while small correspond to the image having larger amounts of a given genetic target. For 3D-SIM, larger intensity values mean that there is more genetic material. Smaller values mean that there is less genetic material. Thus, the max and min are explainable since it describes the least and greatest amount of the genetic target in a given observation, respectively. It is interpretable since max and min values corresponds to physical definitions about the genetic target of interest.

The skewness values, 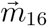, indicate how much the values of the target CD intensity values differ from a symmetric distribution (assuming that the target CD’s distribution of intensity values is unimodal). A value of zero means that the CD’s intensity values are fairly symmetrical, while non-zero values indicate that the distribution has a long tail. Negative values indicate that the left tail is longer, while positive values indicate that the right tail is longer. Thus, negative values indicate that there are outliers of dense areas of genetic material. Positive values indicate that there are outliers of sparse areas of genetic material. Therefore, skewness is explainable since it describes an aspect of the CD’s intensity distribution. It is interpretable since it indicates the amount of genetic material present.

The kurtosis values, 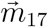, indicate the presence of outliers[49]. Assuming that the intensities follow a normal distribution, a value close to 0 indicates that there are no outliers. Non-zero values indicate the presence of outliers. In other words, a value close to 0 indicates that the CD is has a very consistent amount of genetic density. Non-zero values indicate the presence of sections of the CD that greatly deviate from the typical genetic density. Therefore, skewness is explainable since it describes an aspect of the CD’s intensity distribution. It is interpretable since it indicates the presence of abnormal sections in the CD.

The last two metrics are merely the proportions of the intensity values greater than 24,000 or 30,000. These metrics were only used for 3D-EMISH. Large values indicate that the CD is sparser, while smaller values indicate that the CD is denser. Therefore, these proportions are explainable since it describes the amount of the CD that is sparse. They are interpretable since it indicates the proportion of the CD that is sparse.

#### A.4 Extended Results for 3D-EMISH

The resulting LASSO plot for evaluating the choice in λ for 3D-EMISH is provided in Figure B2. The bottom axis displays the log of the tuning parameter. The y-axis displays the misclassification error for varying tuning parameter values. The top axis displays the number of features retained in the model. The left dotted line displays the optimal λ value. The right dotted line displays the λ within 1 standard error of the optimal one. The red dot is the mean misclassifcation for a given λ. The error bars are the associated values obtained across the 10-folds. The optimal value was estimated to be about 0.01. This was used for all of the 3D-SIM experiments since it is a more prudent choice of λ.

Table C1 provides the numerical values for the relative VI. For example, E1 and E3 are only about 30% as important as the Q3 intensity.

Tables C2 and C3 provide the confusion tables for the training and validation data. Since we have large diagonal values with smaller non-diagonal values, we have evidence that the model performs well on the data used to build the model and data that the model was not trained on.

#### A.5 Extended Results for 3D-SIM

One of the notable choices made for the 3D-SIM models was the λ parameter for all of the models as seen in Figures B3, B4, and B5. These figures have similar interpretations as provided in Figure B2. While the procedure produced different final values (which provides additional, albeit weaker, evidence that the open and closed CD’s physical characteristics are changing). However, across all of the data sets (including 3D-EMISH), the same general shape was provided. Further, the authors of the LASSO have stated that the more appropriate value for the LASSO is the larger value suggested value since it “errs on the side of parsimony”[31]. In other words, assuming that the evaluation metric is “reasonablly” close, choosing the simpler model is preferable since it favors a simpler model. To that end, we selected the most conservative λ value across all of the experiments (about 0.01), as the value for the 3D-SIM models. This value favors a more conservative model that actively avoids over-fitting to the data, while also providing more interpretable and explainable models since less variables are needed to classify open and closed CDs. This choice of a more conservative value for was also motivated by a personal effort to provide a more prudent solution in an effort to temper our model performance.

Tables C4–C6 provides the numerical values for the relative VI for the control, 6 hour treatment, and 30 hour treatment, respectively. Max and Surface Area remain the most important variables across the three, but the number of important variables increases with the length of the treatment.

Tables C7–C12 provide the confusion tables for the training and validation data across the control, 6 hour treatment, and 30 hour treatment cells. Since we have large diagonal values with smaller non-diagonal values for each confusion matrix, we have evidence that the model performs well on the data used to build the model and data that the model was not trained on.

#### A.6 Image Processing for 3D-EMISH

The exact image processing steps differ from the more general framework provided in the main text in Equations 1 - 4. Specifically, we first apply

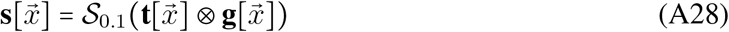

where 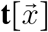 is the input image that contain a chromatin structure (CS), ⊗ is the convolution operator, 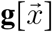 is the local minimum operator of a 2 × 2 × 2 voxel, 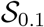 is the Gaussian smoothing operator with a standard deviation of 0.1, and 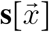 is the resulting smoothed image. We first removed spurious noises using the local minimum using a 2 × 2 × 2 window. We then further smoothed the image using a Gaussian operator to remove the block-like structure created by the previous operation. Examples of the input image and resulting image are provided in Figures B7 and B8, respectively.

Next, we performed

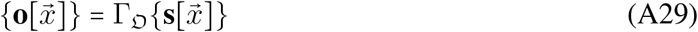

where 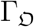 applies Otsu thresholding to each slice of the input smoothed image-stack. This operation extracts the structure from the background without the need for human inputs. An example of this result is provided in Figure B9.

Next, we applied

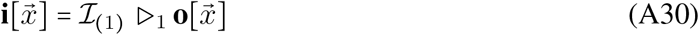

where 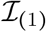 extracts the largest object from the image-stack and 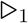 erodes the isolated object 1 times. We first eroded the image isolate spurious aspects of the chromatin structure. We then isolated the largest object from the surrounding noise in the image. An example of this result is provided in Figure B10.

We then applied

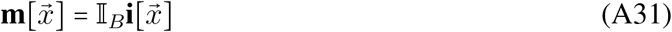

where 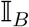 interpolates the object using the other slices to construct the missing slices. We performed this step to provide an estimate of the chromatin physical structure in physical space. An example of this result is provided in Figure B11.

To determine the number of CDs, we then need to apply the mask to the original image. We do this by

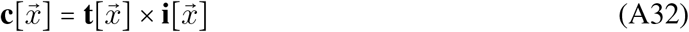

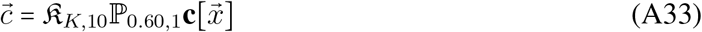

where 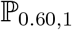 finds the local max peaks of the structure from the input image while ignoring those intensities less than the 60*^th^* percentile of intensity values of the structure while requiring a distance of 1 voxel for each peak. The next image operator, 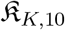 performs *k*-means clustering on the potential peaks to identify the centers of each domain with automatic cluster determination. We consider up to 10 potential clusters. This outputs *k* centers in a vector 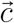. These operators were performed to find the center of each CD in the CFS. We then needed to extract the CDs in the correct units. Thus, we performed:

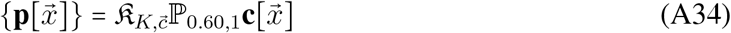

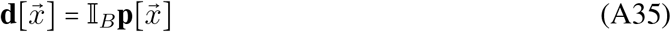

where we first predicted which domain each core of the structure in 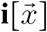 belonged to in Equation A34. We then interpolated in Equation A35 as was done previously in Equation A31. An example of the discovered CDs is provided in Figure B12.

#### A.7 Image Processing for 3D-SIM

There are multiple targets in the colon cancer cell image, this differs from the data obtained from 3D-EMISH data. The 3D-EMISH images had a single target in each image. Thus, some of the image processing steps need to be altered for us to collect the metrics.

Since the 3D-SIM images have three channels, we first need to relevant channels. To that end, we first apply

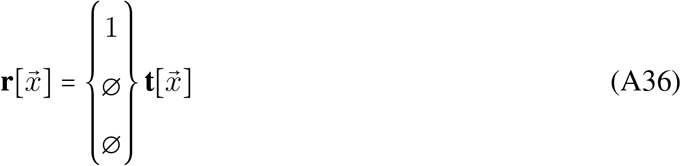

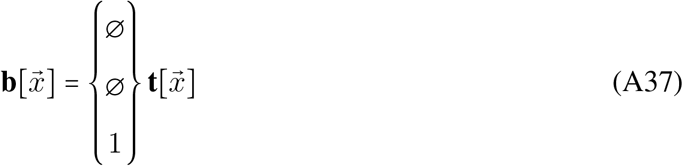

where we extracted the blue channel for the DAPI stained intensities and red channel for H3K27me3 intensities. 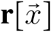 and 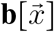 are shown together in Figure 1e. The DAPI channel will be used as a mask to help remove the spurious signals outside of the nucleus. We then applied Equation A28 to smooth 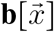 and obtain 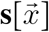 as seen in Figure B13.

We then applied

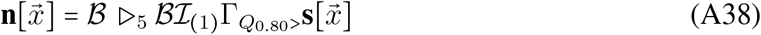

where *Q* is the quantile operator, 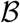 is the binary fill hole operator, and 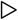 is the erosion operator in order to extract mask of nucleus to extract only relevant H3K27me3 signals. The quantile operator was applied to ensure that only the strongest signals were retrained from the original image. The isolation, binary fill holes, and erosion operators were applied to help identify the nucleus while also filling in the holes missed by the DAPI staining to ensure that every part of the nucleus is extracted. An example is shown in Figure B14.

We then need to extract the relevant H3K27me3 signals. To that end, we apply:

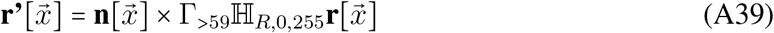

where 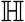 rescales the intensities from 0 to 255 in order to rescale and isolate the relevant signals.

This results in an image of the mask of each CD in a single image as seen in Figure B15.

To extract the relevant image masks for the individual CDs, we then apply

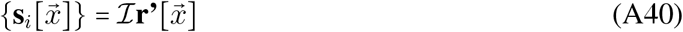

which provides the set of individual CD masks such that *i* ∈ {1, 2, …, *n*}. We then apply Equation A35 to 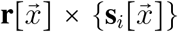 and 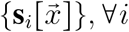 to interpolate the image to ensure that each voxel is approximately the same in the *x* – *y* – *z* directions. Figure B16 provides an example of all the CDs extracted from the example input image.

We then extract the shape and intensity metric using Equation 5. Lastly, we then apply Equations 7 - 10 as in the usual case to classify the open and closed objects. However, we only have one experiment or batch in this case. Conversely, we have three different treatments. Thus, we will build three separate models for each of the three cells: the control, the 6 hour treatment, and the 30 hour treatment.

### B Supplementary Figures

**Figure B1:**
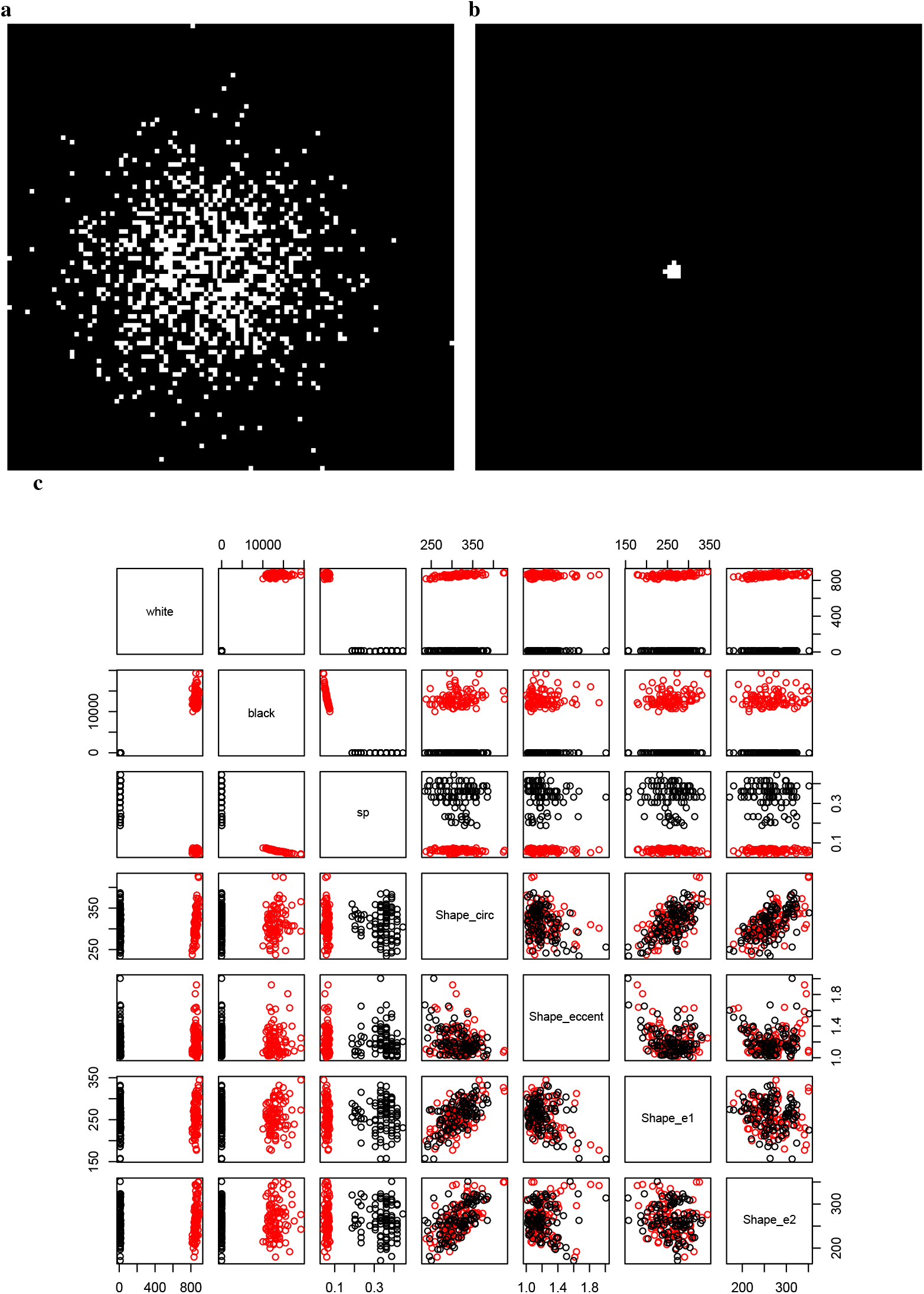
Figure shows examples of the dense and spare simulated genomic structures in Figures B1a and B1b. The resulting shape metrics from 200 simulated structures in Figure B1c shows that the two classes are separable. **(a)** A simulated sparser distribution. **(b)** A simulated dense distribution. **(c)** The collected shape metrics.

**Figure B2:**
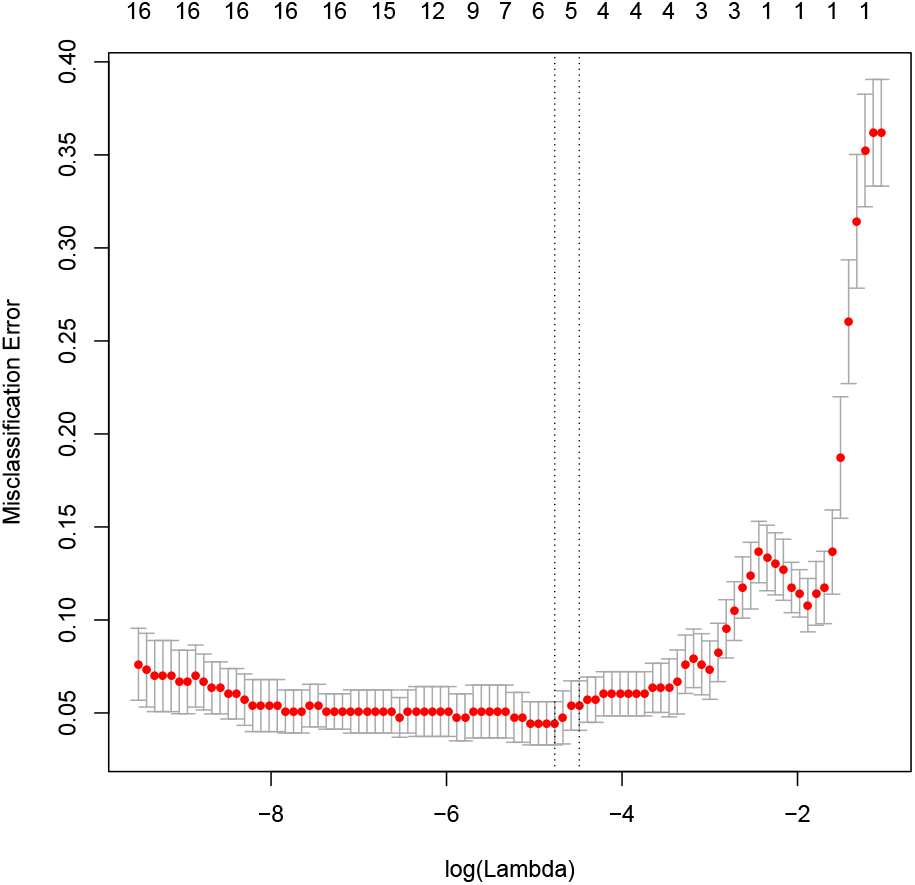
Figure shows the use of 10-folds CV to help select the tuning parameter for the 3D-EMISH data.

**Figure B3:**
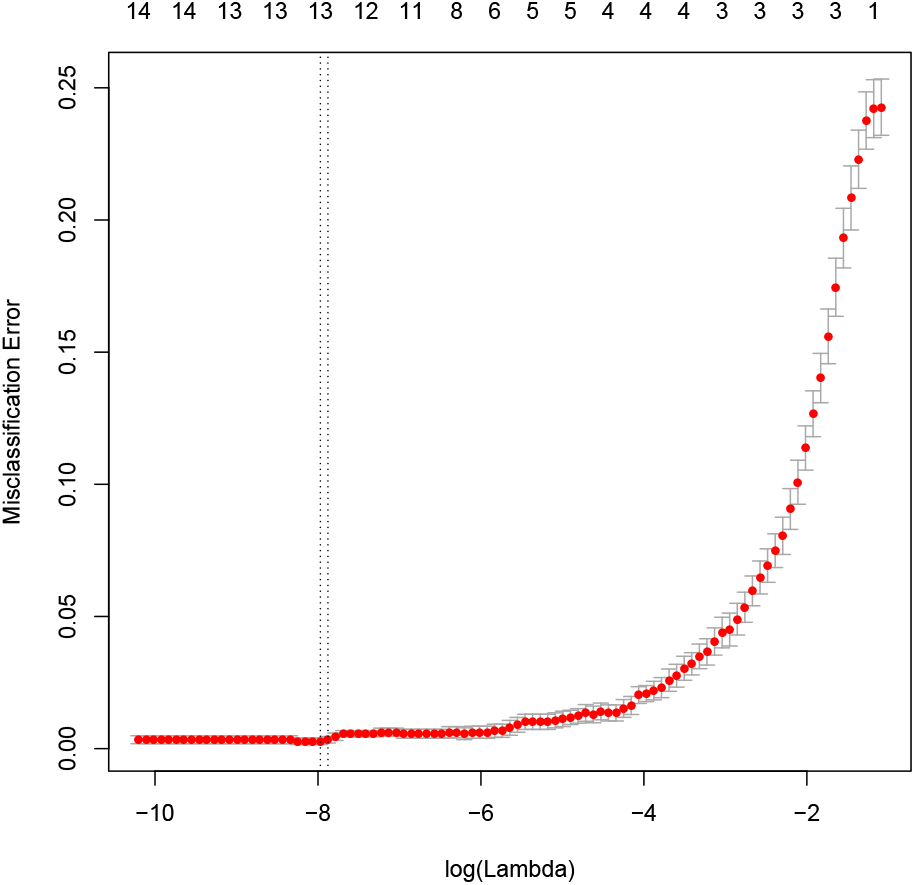
Figure shows the use of 10-folds CV to select the tuning parameter for the control 3D-SIM data.

**Figure B4:**
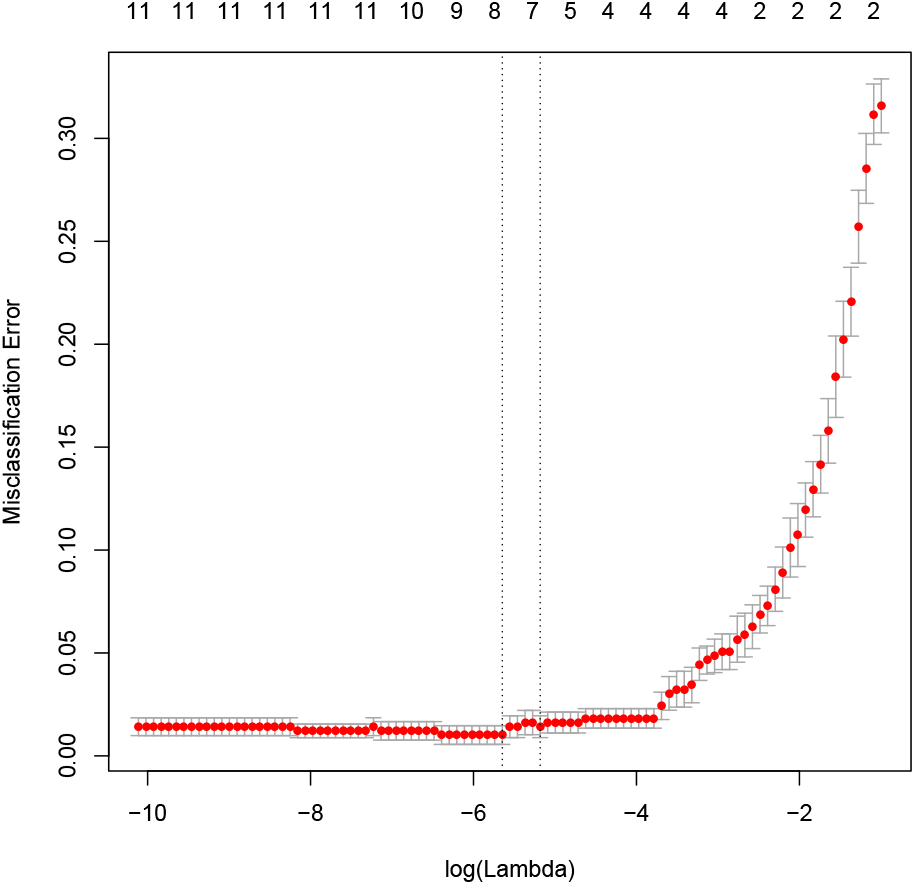
Figure shows the use of 10-folds CV to help select the tuning parameter for the 6-hour treatment 3D-SIM data.

**Figure B5:**
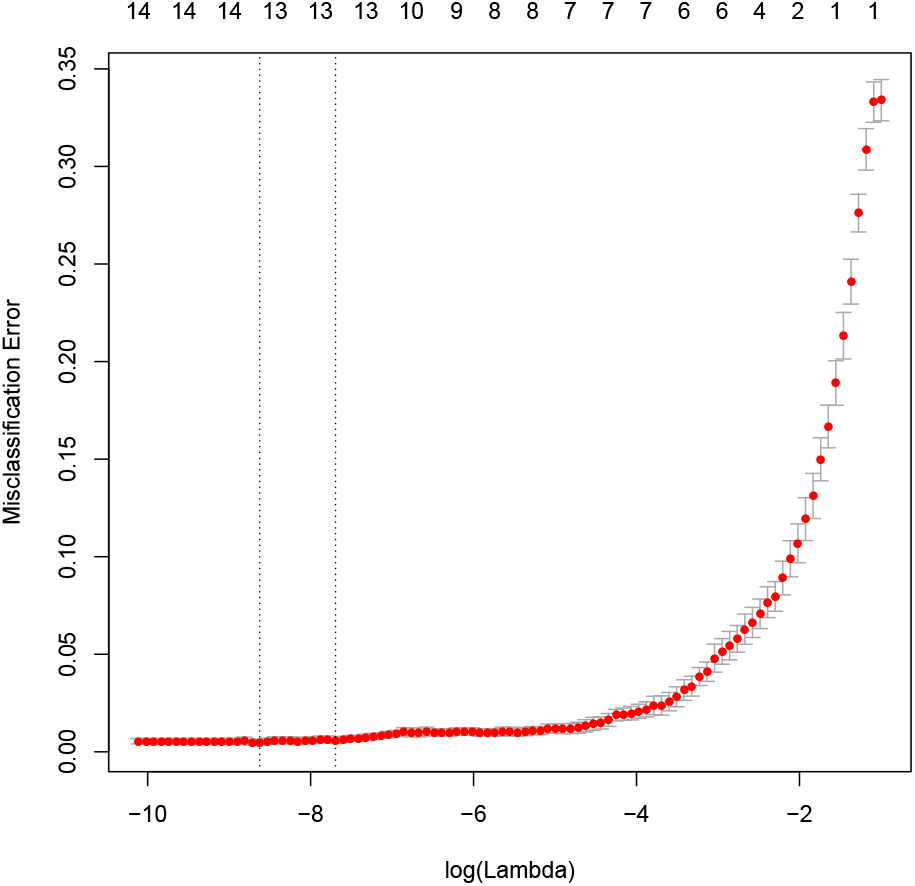
Figure shows the use of 10-folds CV to help select the tuning parameter for the 30-hour 3D-SIM data.

**Figure B6:**
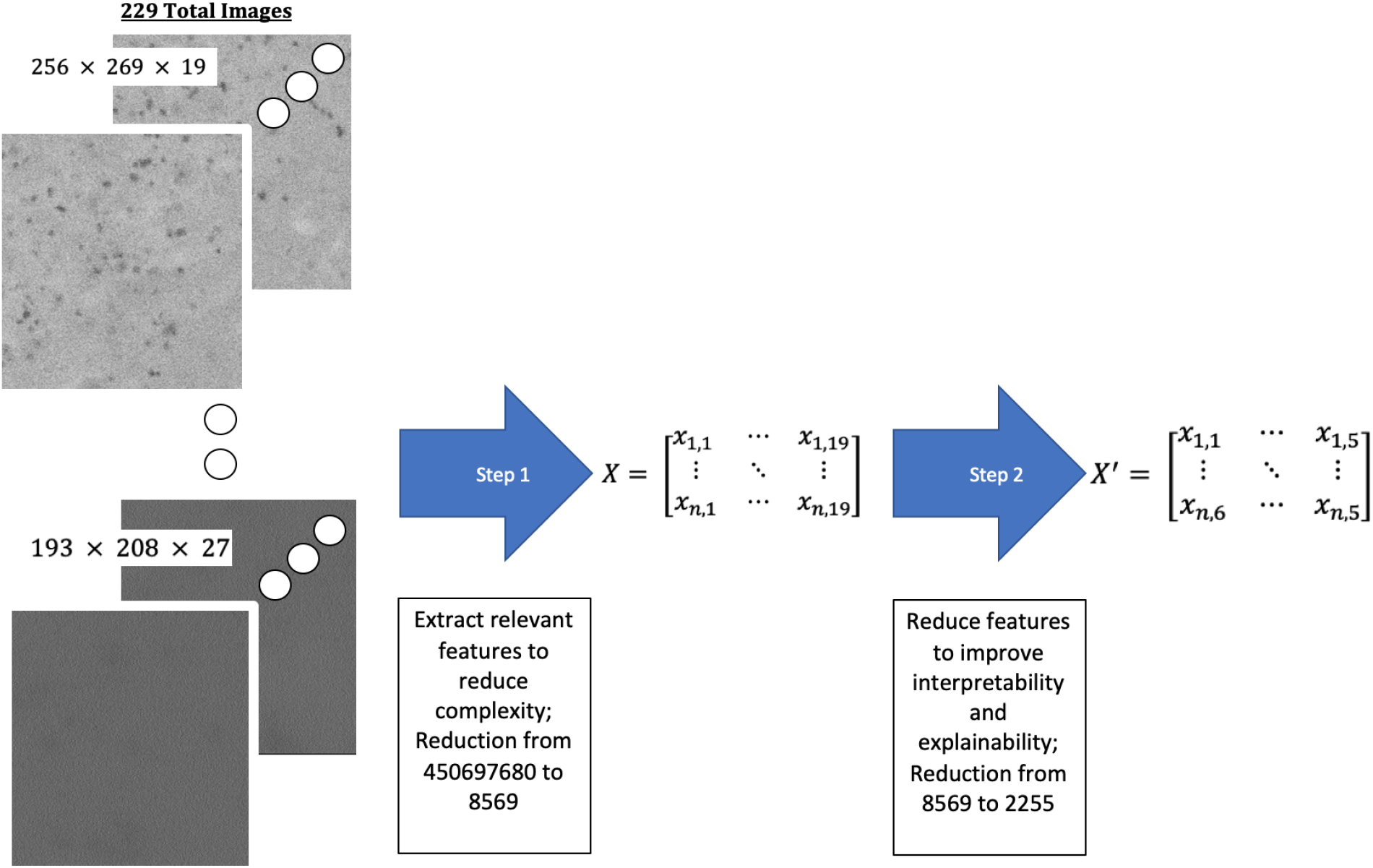
EPICS reduces complexity by removing less important variables by extracting shape and intensity image features and then using the LASSO ML algorithm as exemplified with the 3D-EMISH data.

**Figure B7:**
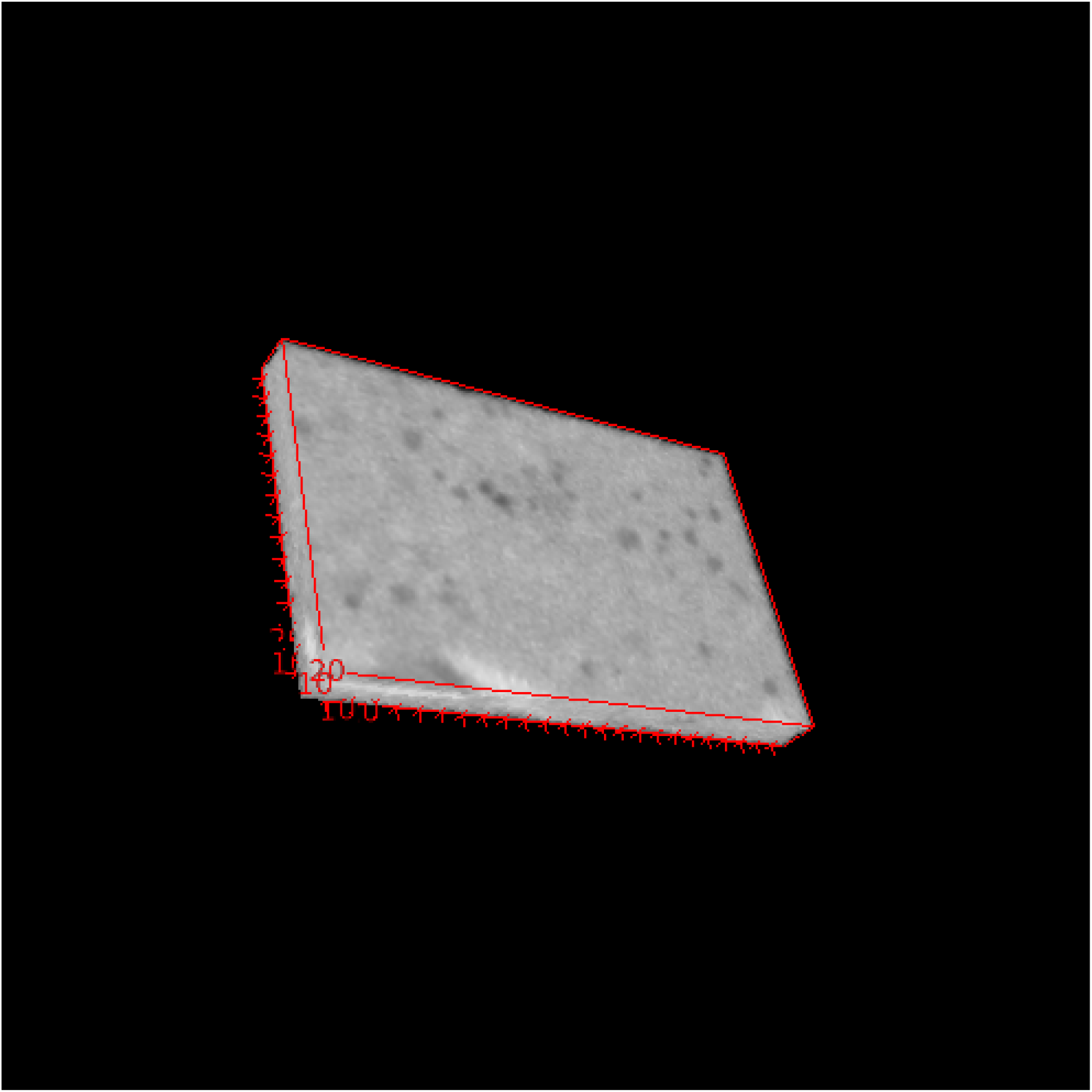
Example of input image for 3D-EMISH.

**Figure B8:**
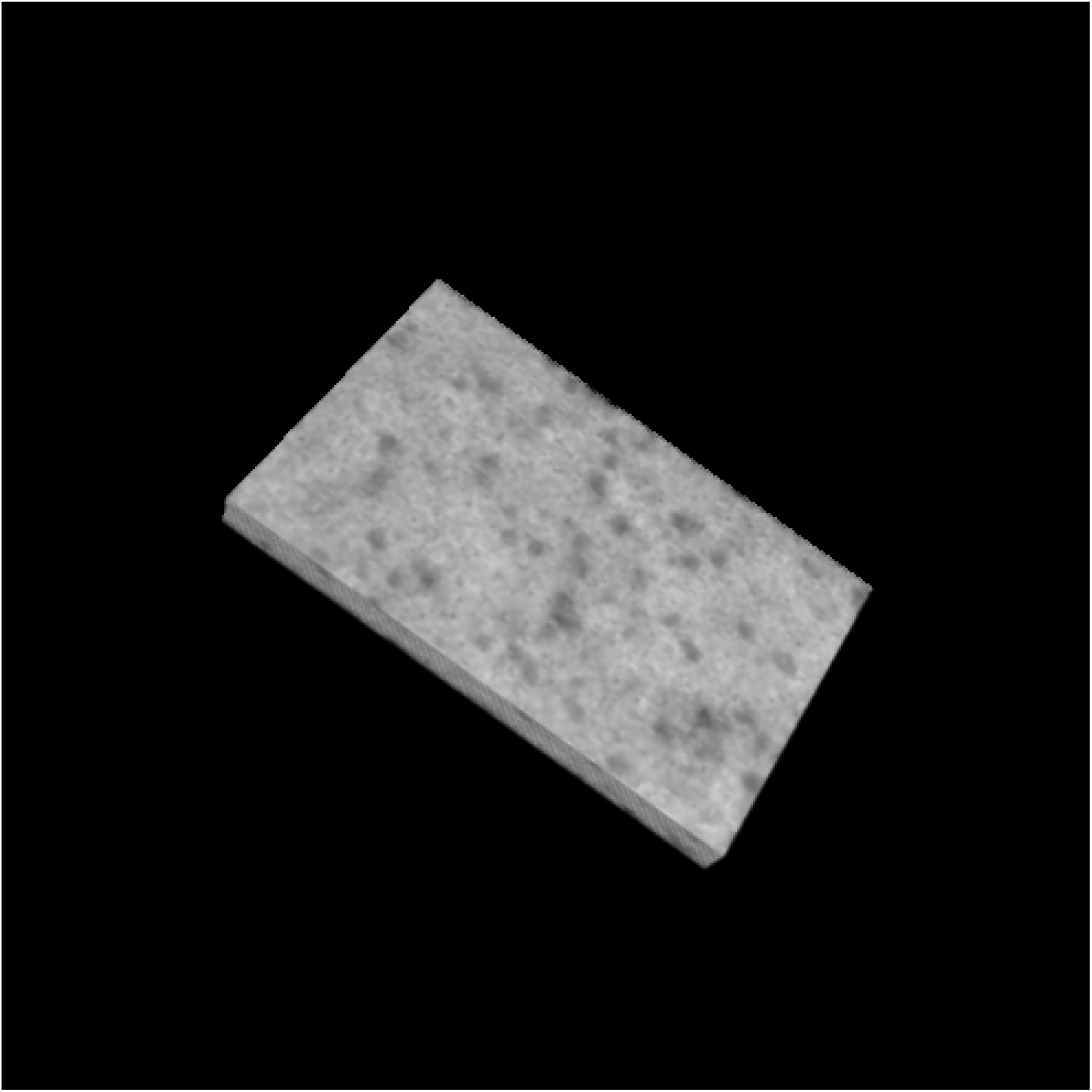
Example of smoothed 3D-EMISH image.

**Figure B9:**
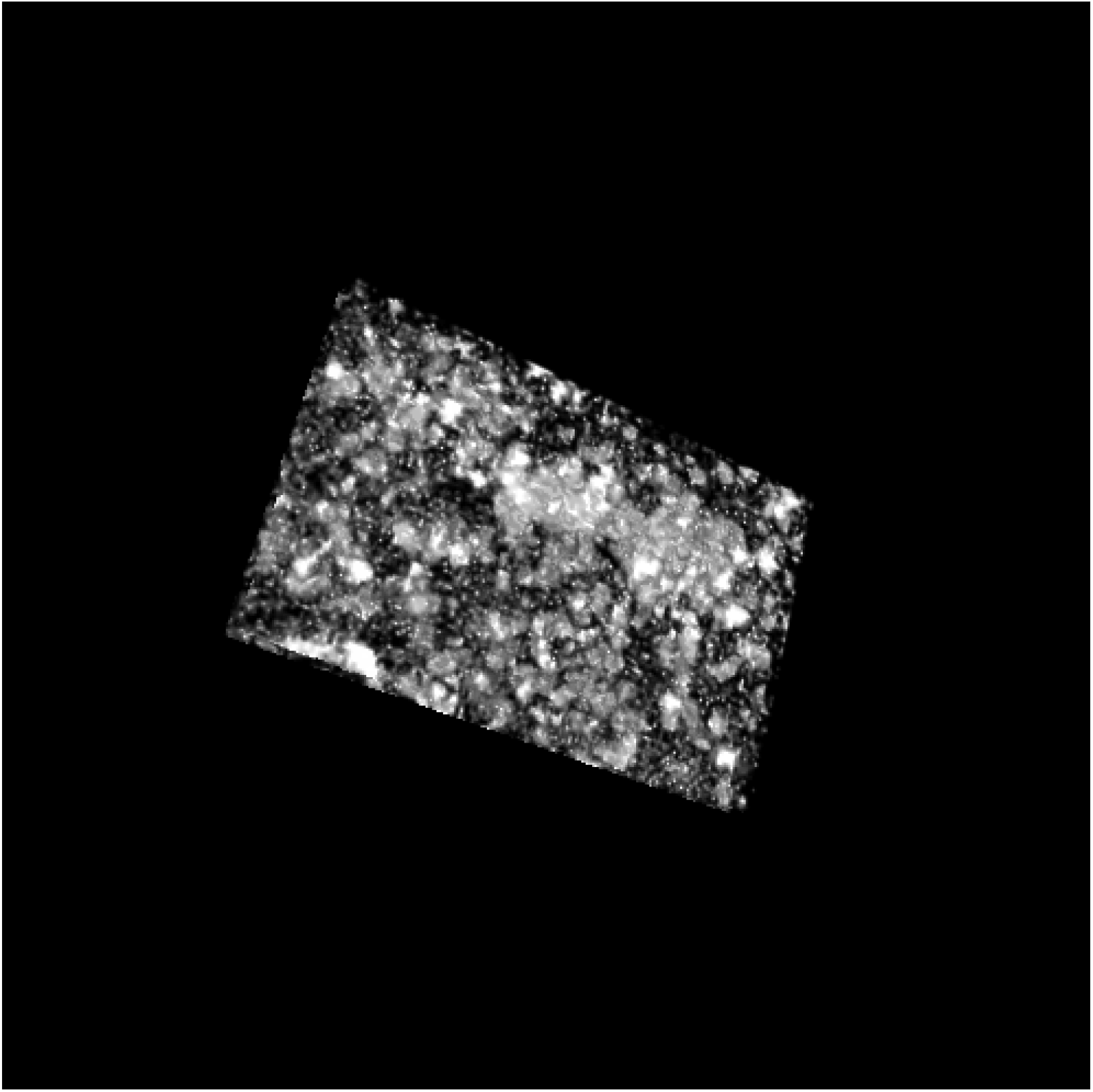
Example of isolated signals from 3D-EMISH example image.

**Figure B10:**
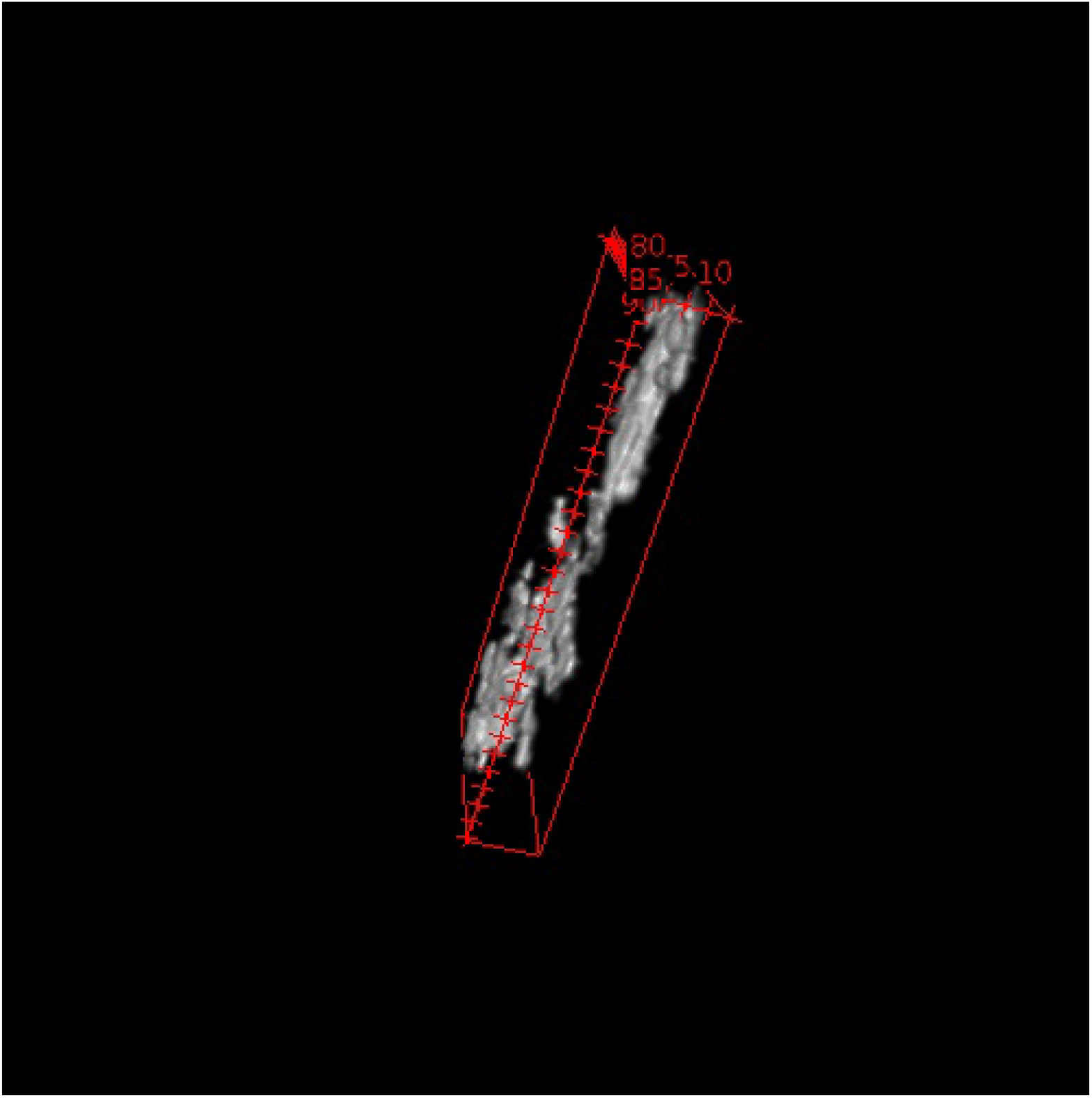
Example of largest isolated object from 3D-EMISH example image.

**Figure B11:**
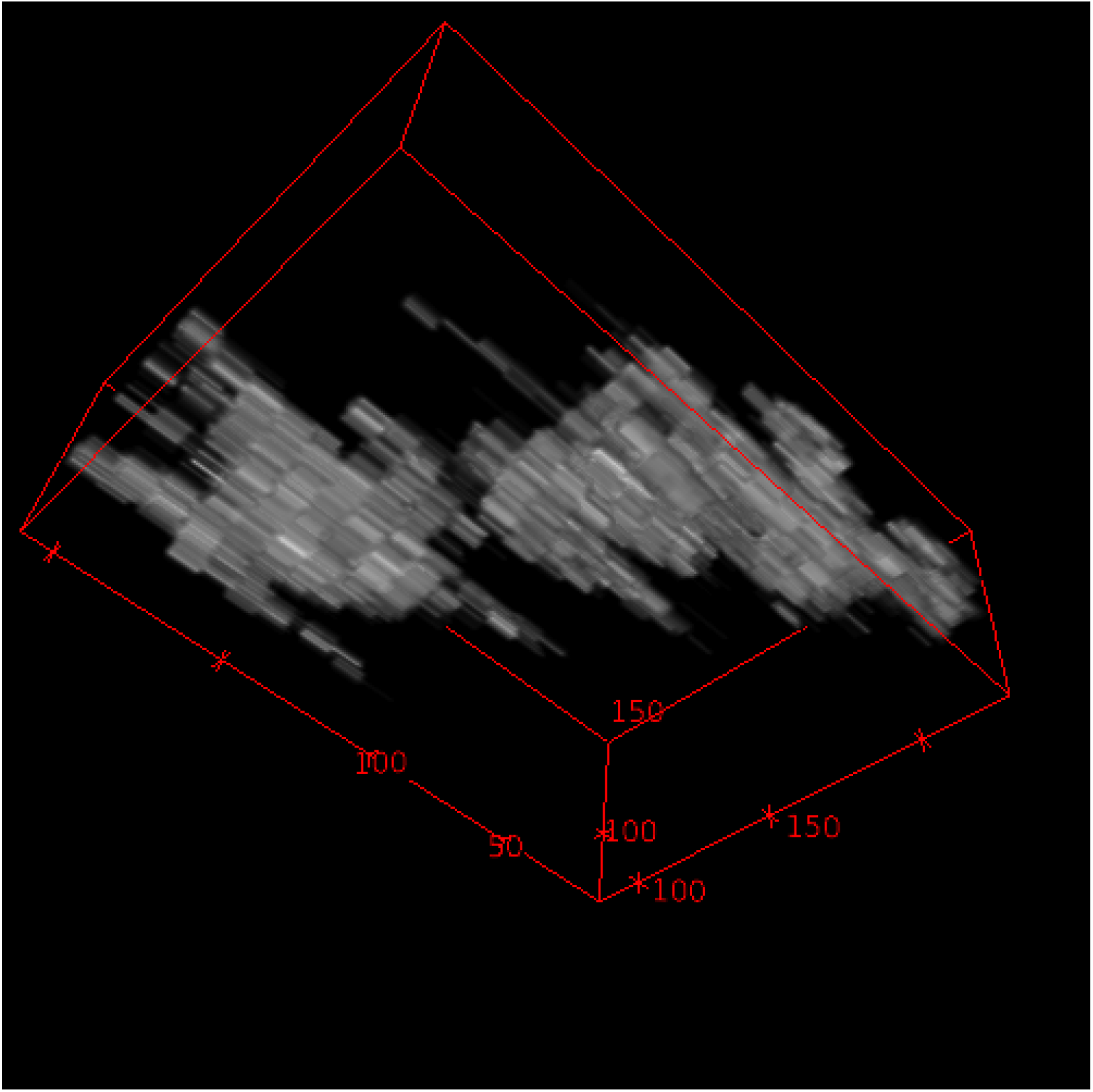
Example of reconstructed object of interest from 3D-EMISH data.

**Figure B12:**
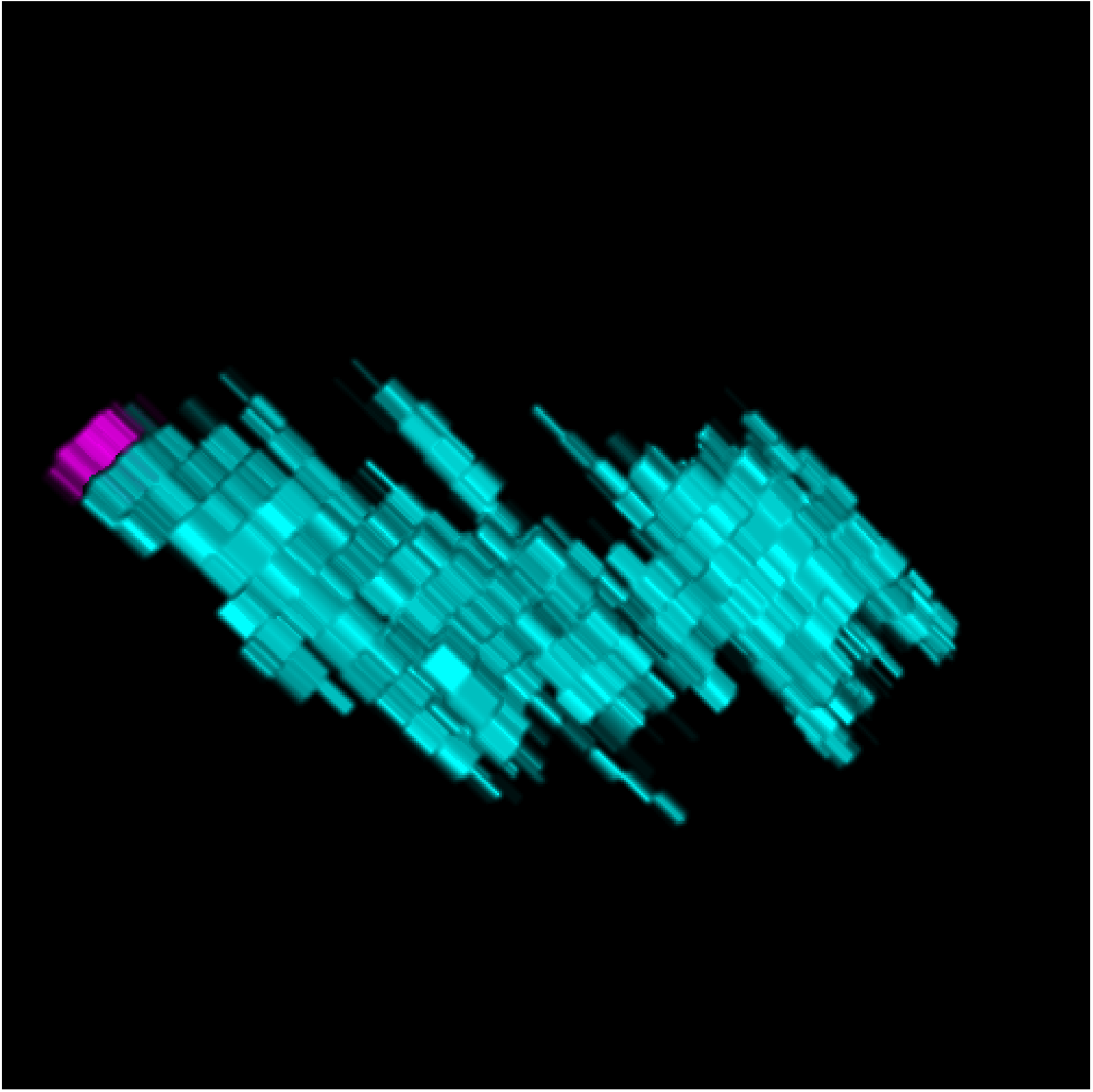
Example of CDs identified from example input 3D-EMISH data. Each different color represents a unique CD.

**Figure B13:**
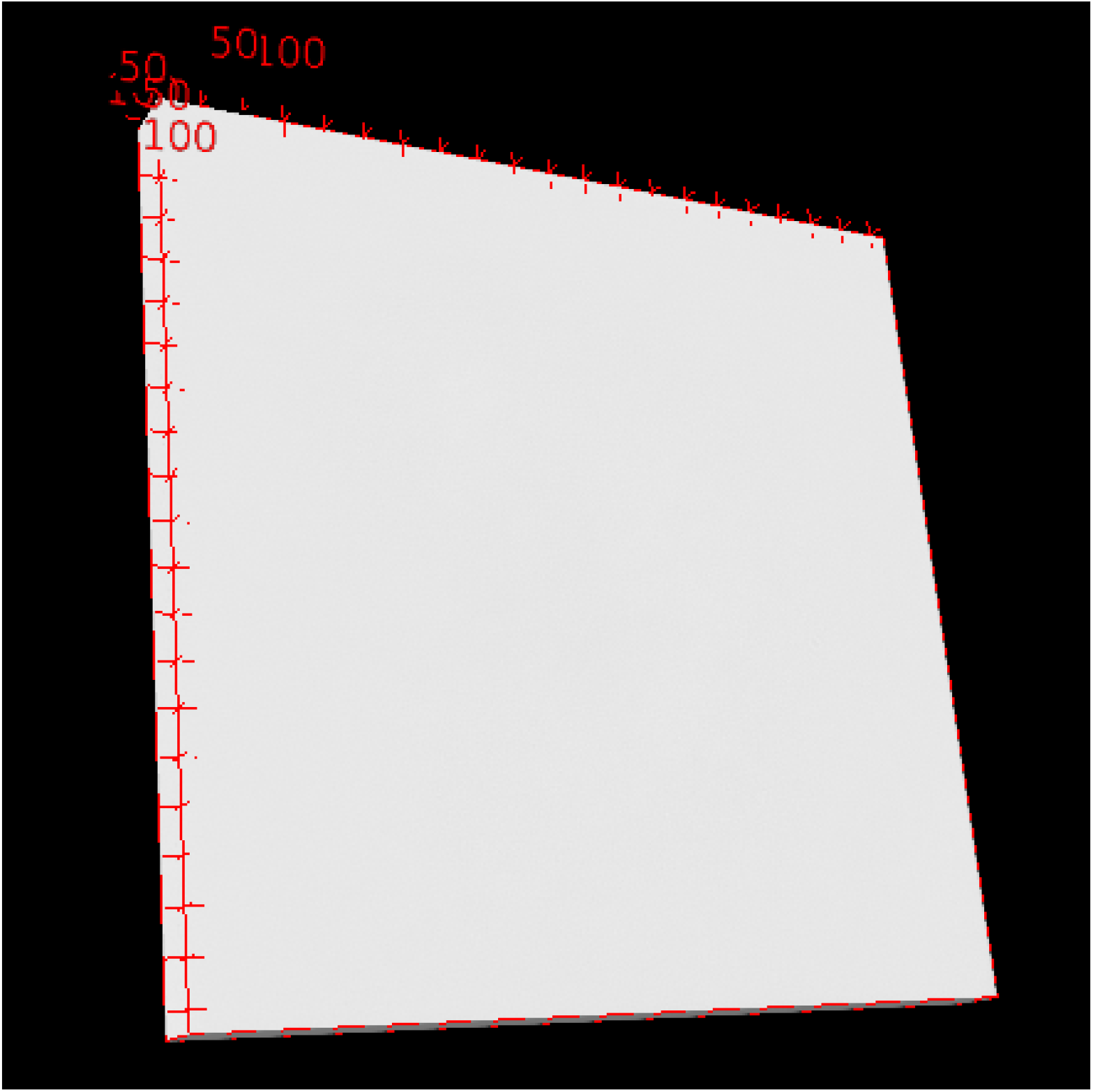
Smoothed image of the DAPI stained image from the 3D-SIM example. The input image was the example image from Figure 1e.

**Figure B14:**
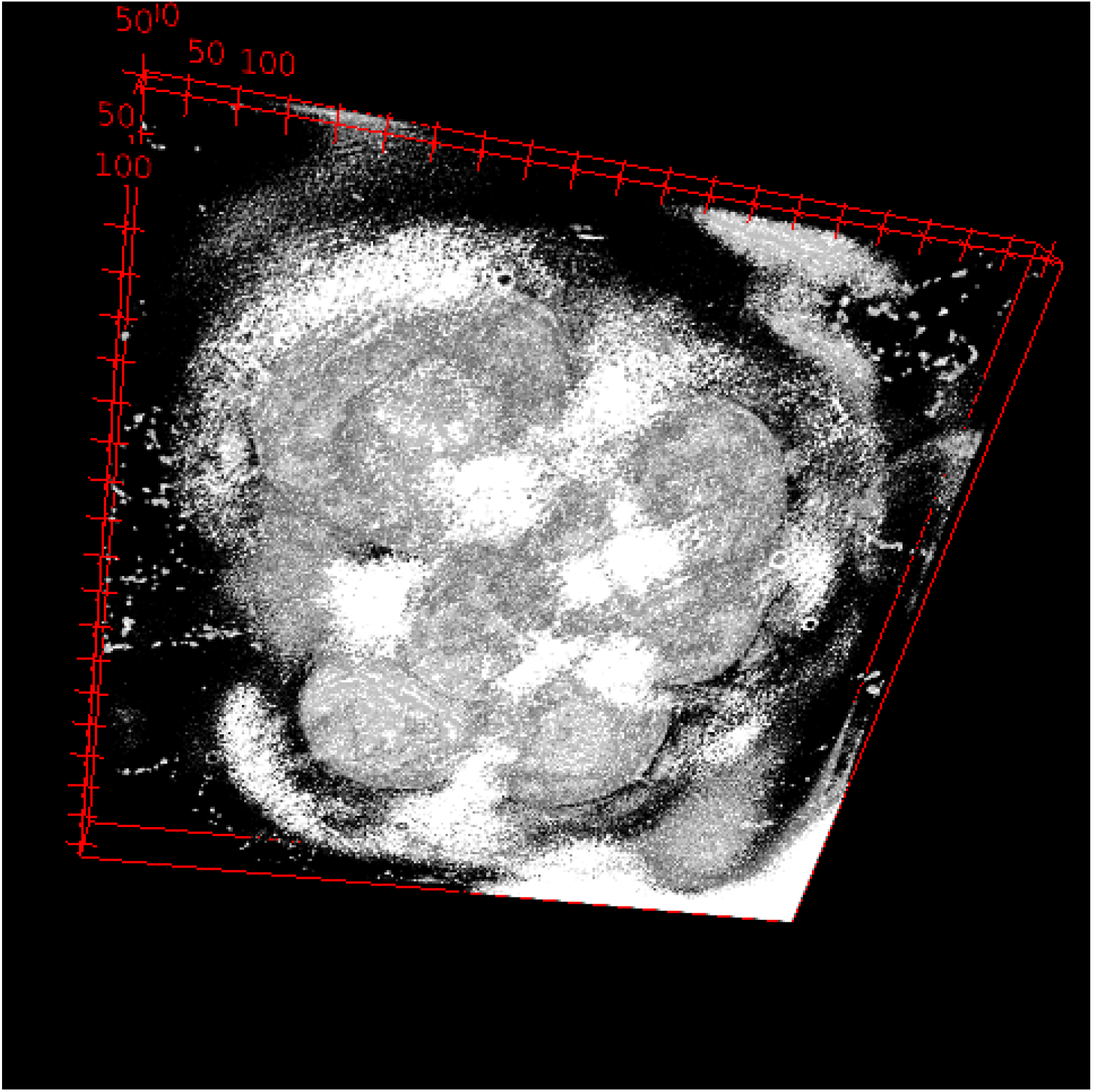
Example of identified mask from the DAPI stained image.

**Figure B15:**
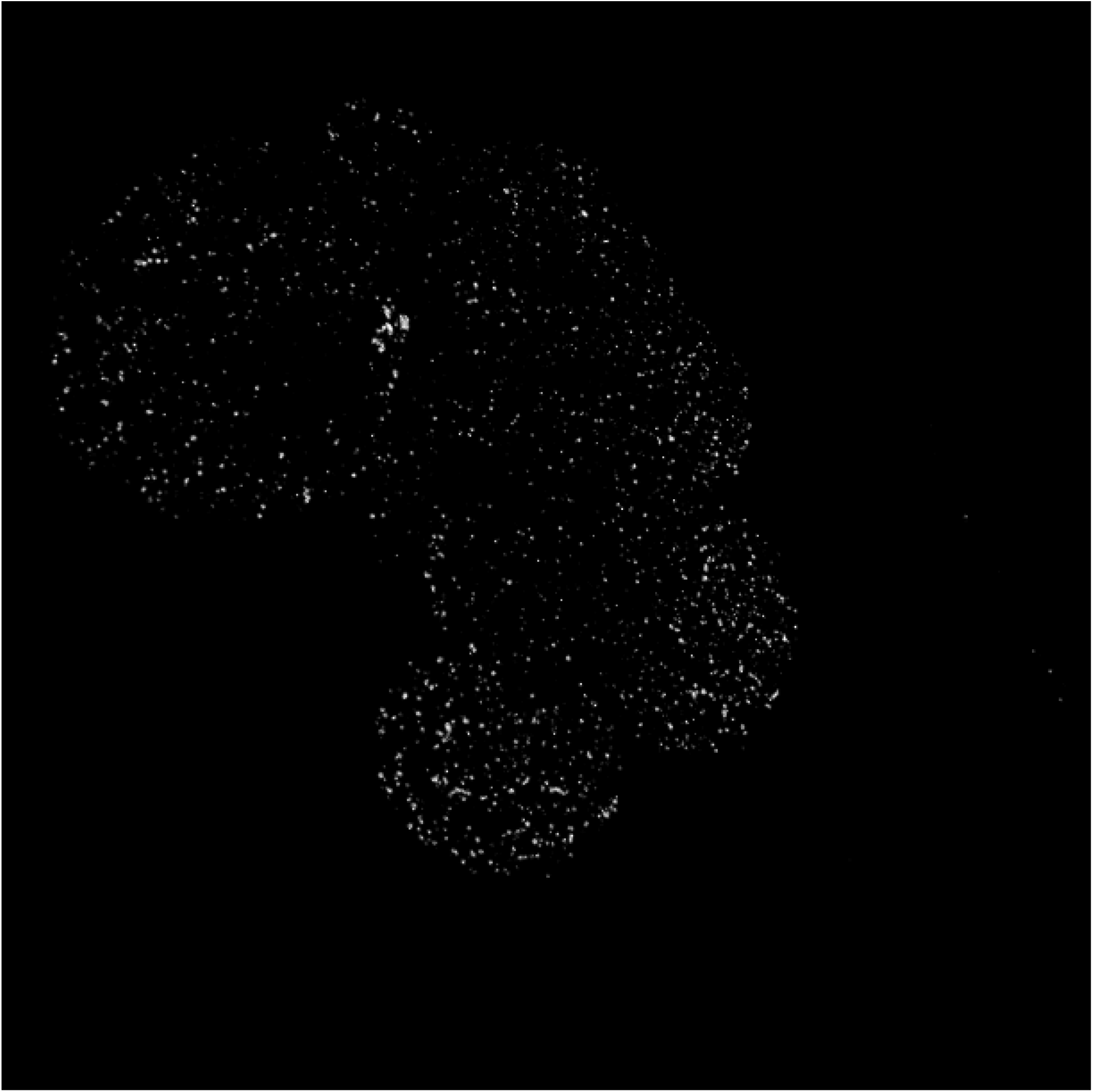
Example of the extracted signals of interest from the 3D-SIM image only within the nucleus.

**Figure B16:**
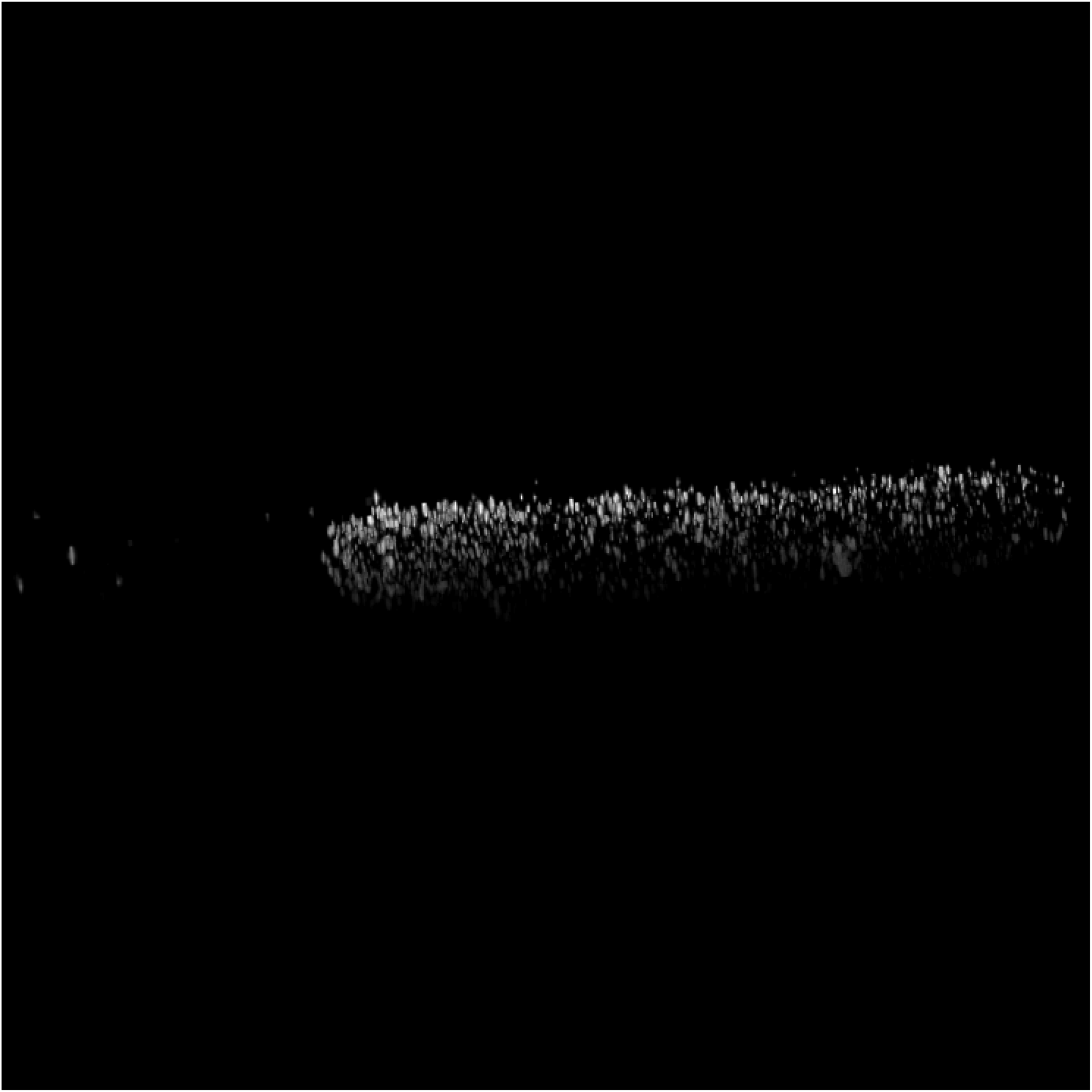
All of the CDs extracted from the example input 3D-SIM image. Each different grayscale intensity represents a unique CD.

### C Supplementary Tables

**Table C1:**
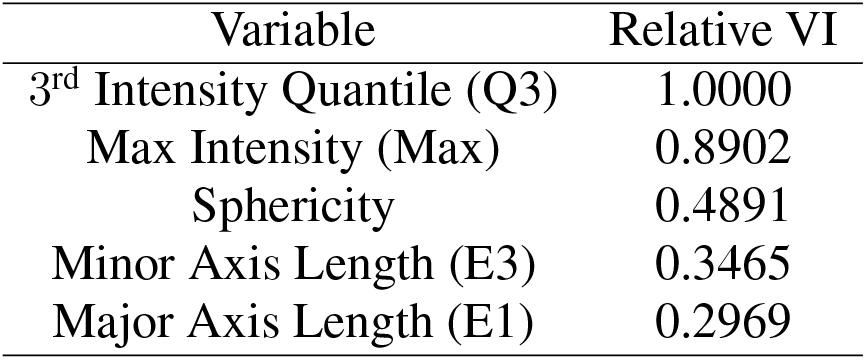
Table shows the relative variable importance (VI) of the classification model for open and closed CDs from the 3D-EMISH data.

**Table C2:**
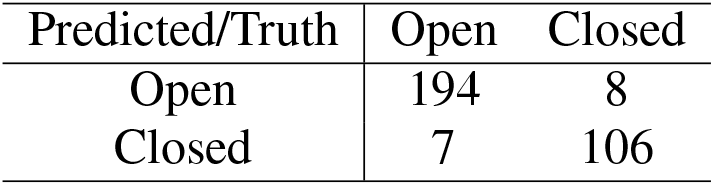
Confusion table on the training 3D-EMISH data.

| Predicted/Truth | Open | Closed |
| --- | --- | --- |
| Open | 194 | 8 |
| Closed | 7 | 106 |

**Table C3:** Confusion table on the validation 3D-EMISH data.

| Predicted/Truth | Open | Closed |
| --- | --- | --- |
| Open | 83 | 2 |
| Closed | 4 | 47 |

**Table C4:**
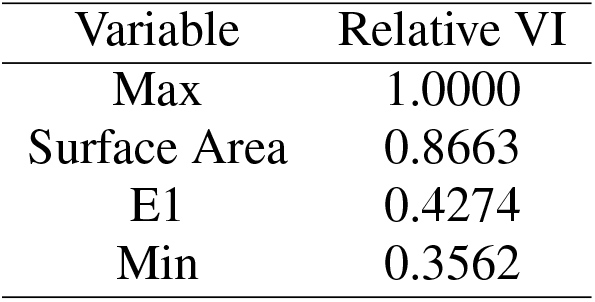
Table shows the relative variable importance (VI) of the classification model for open and closed CDs from the control 3D-SIM data.

| Variable | Relative VI |
| --- | --- |
| Max | 1.0000 |
| Surface Area | 0.8663 |
| E1 | 0.4274 |
| Min | 0.3562 |

**Table C5:** Table shows the relative variable importance (VI) of the classification model for open and closed CDs from the 6 hour 3D-SIM data.

| Variable | Relative VI |
| --- | --- |
| Surface Area | 1.0000 |
| Max | 0.7828 |
| Skew | 0.7127 |
| SD | 0.2293 |
| E1 | 0.1908 |

**Table C6:** Table shows the relative variable importance (VI) of the classification model for open and closed CDs from the 30 hour 3D-SIM data.

| Variable | Relative VI |
| --- | --- |
| Surface Area | 1.0000 |
| Max | 0.9474 |
| Skew | 0.8662 |
| SD | 0.5919 |
| Min | 0.5783 |
| E1 | 0.4860 |
| E3 | 0.3890 |

**Table C7:**
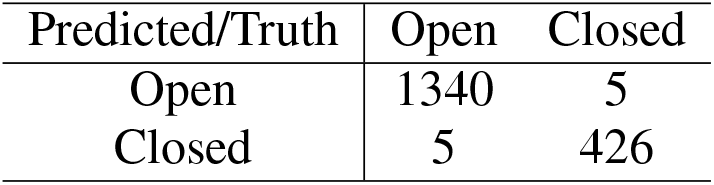
Confusion table on the training data for the control 3D-SIM data.

| Predicted/Truth | Open | Closed |
| --- | --- | --- |
| Open | 1340 | 5 |
| Closed | 5 | 426 |

**Table C8:** Confusion table on the validation data for the control for the 3D-SIM data.

| Predicted/Truth | Open | Closed |
| --- | --- | --- |
| Open | 573 | 2 |
| Closed | 4 | 184 |

**Table C9:** Confusion table on the training data for the 6-hour treatment for the 3D-SIM data.

| Predicted/Truth | Open | Closed |
| --- | --- | --- |
| Open | 338 | 0 |
| Closed | 0 | 156 |

**Table C10:** Confusion table on the validation data for the 6-hour treatment for the 3D-SIM data.

| Predicted/Truth | Open | Closed |
| --- | --- | --- |
| Open | 146 | 0 |
| Closed | 0 | 68 |

**Table C11:** Confusion table on the training data for the 30-hour treatment for the 3D-SIM data.

| Predicted/Truth | Open | Closed |
| --- | --- | --- |
| Open | 1337 | 9 |
| Closed | 9 | 666 |

**Table C12:** Confusion table on the validation data for the 30-hour treatment for the 3D-SIM data.

| Predicted/Truth | Open | Closed |
| --- | --- | --- |
| Open | 574 | 5 |
| Closed | 4 | 285 |

## Notes

https://github.com/zang-lab/epics

### Competing Interest Statement

The authors have declared no competing interest.

